# Neural circuits regulating social dominance implement a strategy predicted by evolutionary game theory

**DOI:** 10.64898/2026.02.20.706896

**Authors:** Donovan Ventimiglia, Elizabeth Chamiec-Case, Claire S. Lee, Bruce Ruff, Kenta Asahina

## Abstract

Social conflict is a fundamental challenge for all animals and determines access to critical resources like mates and food. Evolutionary game theory predicts that natural selection should yield competitive strategies that balance the benefits and costs of social conflict. However, whether such strategies are embedded within the neural circuits that regulate aggression remains unclear. Here, we identify a neural circuit regulating the decision to flee during fighting in male *Drosophila* and show that the onset of defeat is governed by a probabilistic strategy predicted by evolutionary game theory. This mechanism arises from the inhibition of Tk-GAL4^FruM^ neurons that promote aggressive arousal in males. Inhibition is mediated by a mushroom body circuit involving PPL1 dopaminergic neurons and V2 mushroom body output neurons, both classically associated with aversive learning. Silencing this circuit disrupts the onset of defeat, while activating it induces rapid defeat. Conversely, activation of reward-encoding PAM dopaminergic neurons promotes winning, revealing a dual role for dopamine in shaping contest dynamics. Finally, we find that internal state variables such as hunger and motivation shift the defeat onset probability distribution, consistent with game theory predictions of how payoff modulates fighting persistence. Together, our results provide direct evidence that evolutionary strategies based on payoff, long described by game theory, are implemented as circuit-level computations that regulate aggression.

## INTRODUCTION

Social conflict is an inherent feature of animal life. Animals fight for food, territory, and mates, and competition for these limited resources plays a central role in animal evolution^1^. While the benefit of winning a fight appears obvious, competition comes with substantial costs including energy expenditure, injury risk, and time diverted from other fitness-enhancing activities^2,3^. This suggests that natural selection should favor the evolution of competitive strategies that balance the benefits of winning with the cost of competition (termed payoff).

Evolutionary game theory supports this idea quantitively^2^, showing that competitive strategies can undergo selection in a population through enhancing individuals’ fitness, becoming evolutionarily stable. However, whether specific strategies through evolution have become embedded in the neural circuits that regulate social conflict, for example in aggression circuits, is unclear. Persistence-based contests are widespread throughout the animal kingdom^1^ and include behavioral tactics such as aggressive displays, agonistic vocalizations, or non-injurious wrestling, all of which can also gate additional forms of aggressive behaviors^2,3^. In these forms of contests, animals must strategically choose how long to participate before conceding and seeking other opportunities. In evolutionary game theory, War of Attrition (WOA) is a model that applies to these types of scenarios, in which individuals do not inflict injuries but instead compete through persisting in costly behaviors until one concedes^2,3^. Under these conditions, game theory predicts that no pure strategy governing the choice to concede (such as always persisting for a set amount of time) is evolutionarily stable. Instead, it is predicted that individuals should use mixed strategies that effectively prevent opponents from predicting persistence-times. Following the assumption that cost linearly increases with time, the evolutionarily stable strategy is to select a contest duration according to an exponential probability distribution that is shaped by the resulting payoff of contest (eq.1)^2^.

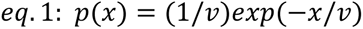

War of Attrition model (WOA): *v* denotes the payoff, defined as the difference in a resource’s value and the cost of persisting in the contest.

This suggests that neural circuits regulating the choice to flee under these conditions may be regulated through a probabilistic selection of persistence-times sampled from an exponential distribution, the shape of which is modulated by payoff.

The common fruit fly *Drosophila melanogaster* offers an opportunity to investigate the neural circuits regulating the critical choice whether to continue fighting or to flee, with the ability to combine quantitative behavioral analysis^4^ and mechanistic dissection of neural circuits with single-cell resolution^5^. *Drosophila* lack weaponry, and fights between males consist of wrestling-like behaviors^6,7^ that align well with the criteria of persistence-based contests. Importantly, fights among *Drosophila* males also result in clear dominance relationships: winners continue to be aggressive and initiate attacks, while losers signifying defeat by halting aggression and fleeing from their opponent^8–11^. Importantly, neural circuits regulating this key decision in losers is not known.

Several distinct neural populations that promote aggressive behavior in *Drosophila* have been identified. These neurons often belong to sexually dimorphic neural circuits that express sex-determining factors *fruitless* (*fru*) or *doublesex* (*dsx*)^12,13^. Of these cell types, the Tk-GAL4^FruM^ neurons are critical regulators of male-specific aggressive arousal levels^14^. Excitation of the Tk-GAL4^FruM^ neurons robustly promotes fighting in males, while silencing of them largely eliminates aggression^10,14,15^. Although the functional organization among aggression-controlling neurons is not fully delineated, evidence suggests that Tk-GAL4^FruM^ neurons are a hub for other aggression-controlling neuronal populations^15–17^. Suppression of this neuronal group could be a well-suited mechanism for controlling the onset of defeat during aggressive contests. However, little is known about how aggression-promoting circuits or neuromodulators contribute to regulating the transition from aggression to fleeing, and consequently, winner-loser dominance formation in *Drosophila*.

Here, we combine behavior, optogenetics and functional calcium imaging to dissect the neural circuits regulating defeat onset. Strikingly, we find that defeat onset in both spontaneous and optogenetically induced fights follows an exponential probability distribution, consistent with the prediction from evolutionary game theory. We show that this probability distribution occurs through the inhibition of Tk-GAL4^FruM^ neurons, which requires PPL1 dopaminergic neurons (DANs) and V2 mushroom body output neurons (MBONs). We show that resource need and arousal levels, as well as V2 MBONs activity levels, modulates this exponential probability distribution, consistent with the role of modulating payoff in the WOA model (eq.1). Our results provide the first glimpse into a circuit-level mechanism that actuates a defined behavioral strategy regulating aggression and supports that natural selection has embedded game-theoretic principles into the neural circuit architectures regulating social conflict.

## RESULTS

### Defeat onset follows game-theory

During fly fights, male flies exchange lunges (a form of physical attack)^6,7^ until the loser concedes by switching to fleeing. Game theory predicts that, when equivalently matched opponents compete, the timing of defeat onset (withdrawal) from the contest should follow an exponential distribution (eq.1). To ask if defeat onset in *Drosophila* matches this theoretical prediction, we recorded a large number of fights under stereotyped conditions and measured the duration of fighting before loser flies became defeated. We first analyzed fighting between age-matched socially isolated male flies in custom-designed fighting chambers, and found that most pairs formed clear winner-loser dominance (Fig. 1a, b; Supp. Fig. S1a). Winners remained aggressive throughout the assay, producing a consistent rate of lunging attacks and chasing their opponent. In contrast, eventual losers exhibited an abrupt behavioral shift in which they would acutely stop attacking and switch to fleeing, which we could observe quantitively by measuring lunging and facing-angle orientation (Fig. 1a, c, d; Supplemental Video 1). Following this sharp transition, we observed that almost all losers (92%) would remain non-aggressive (Fig. 1e), consistent with previous observations suggesting that defeat triggers a persistent change in internal state^8–10,18^. This provided an unambiguous marker of when losers chose to exit the competition. We determined the moment of defeat by quantifying when losers ceased lunging and their facing angle orientation shifted away from their opponent (see Methods). Using this transition point, we then measured the time duration before losers succumbed to defeat (defeat onset) in fly-pairs which both flies displayed active fighting (lunging).

**Figure 1.**
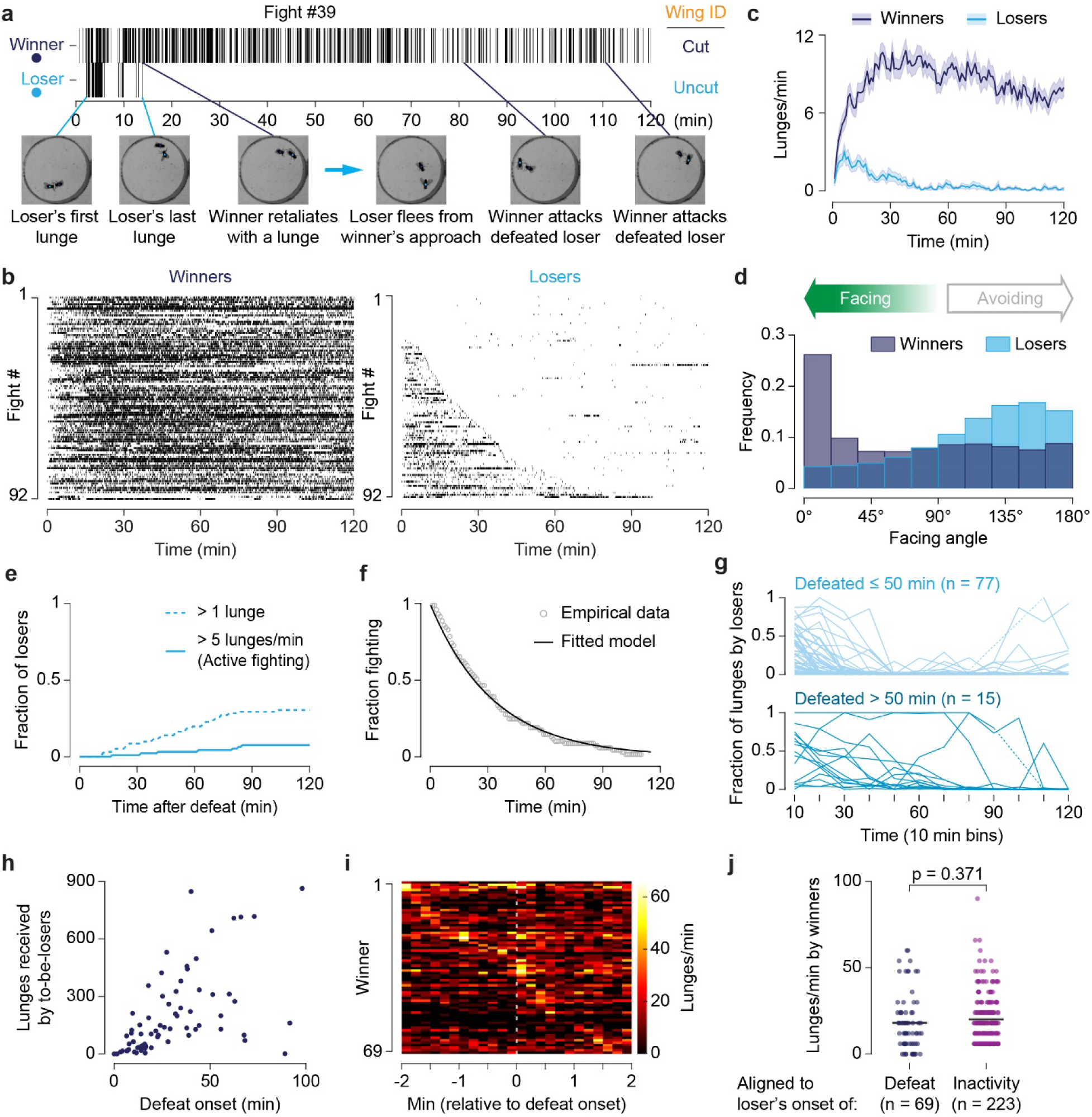
Defeat onset time during dominance formation follows WOA game-theory predictions. **a)** Raster plot of lunges exchanged between an example pair of flies that formed a winner-loser hierarchy. Top: Lunge raster. Bottom: Representative frames from stereotypical fighting phases that occurred before defeat. **b)** Paired lunging rasters of socially isolated wild-type (Canton-S) flies sorted by defeat time, showing behavior of winners (left) and their paired losers (right). Each row corresponds to a single fly; pairs are aligned across panels by fight number. Ticks indicate individual lunges. **c)** Mean lunge rates (1-min bins) of winners (dark blue) and losers (cyan). Trace shading = S.E.M. Data from b (n = 92). **d)** Normalized histograms of facing angles in the 20-minute period following social defeat. Data from (b) (n = 92). **e)** Cumulative fraction of loser flies that recovered from defeat and started attacking again. Less than 8% of defeated flies re-engaged in active fighting (>5 lunges/min) within the assay duration. A 2-min sliding window was used for each cumulative count category. **f)** Dynamics of defeat onset. Proportion of eventual losers engaged in fighting over time (dots, 1-min bins), fit by an exponential function (τ = 33.22 min, 95% CI = 32.60–33.84). Loser flies that didn’t initially fight back or recovered from defeat were excluded from the dataset shown in (b) (see methods), n = 68 losers. **g)** The fraction of lunges performed by the eventual loser over time (number of lunges by loser / total lunges per pair). Top plot, flies defeated within 50 min after the onset; bottom plot defeated after 50 min. Dotted lines connect missing time-bins where neither fly lunged. **h)** Total number of lunges received by each loser prior to defeat, plotted against their defeat onset time, n = 92. **i)** Heatmap plot of winners’ peak lunge rate (10s bin) aligned to their opponents defeat onset time (t=0). Each row is a winner fly, sorted by time of peak lunge rate during the 4-min window. Pairs in which the loser was defeated within the first 2 min were excluded, n = 69 winners. **j)** Peak winner lunge rates upon loser defeat (±10-s bin) (n = 69, as in Fig. 1i) or pre-defeat inactivity bouts when the loser did not lunge for at least 1 min (223 bouts, 58 flies). Welch’s unpaired t-test.

Consistent with WOA predictions, contest durations followed a clear exponential distribution (Fig. 1f). Despite near-identical genetics and experimental conditions, individuals varied dramatically in their contest duration. Within the WOA framework, the contest outcome is governed by the minimum of the two persistence times, implying that defeat onset reflects the loser’s withdrawal threshold rather than discrete actions imposed by the winner. Consistent with this, we identified many instances where the dominant attacker would abruptly switch into a defeated state to become the eventual loser (Fig. 1g). Losers also varied dramatically in the number of attacks they received prior to defeat: some received only a few lunges, while others sustained hundreds of attacks (Fig. 1h). Furthermore, defeat onset did not correlate with peak lunge rates in the winner (Fig. 1i), and winner lunge rates upon defeat were comparable to lunge rates observed during other periods of loser inactivity prior to defeat (Fig 1j, Supp. Fig. S1b,c). Together, these patterns are consistent with the WOA game-theory model, in which the action to flee is primarily generated from a probabilistic selection of persistence-time.

### Defeat onset suppresses Tk-GAL4^FruM^ neurons

We next asked whether these defeat dynamics reflect the action of a dedicated neural circuit. If a specific neuronal mechanism acts to trigger defeat and suppress aggression, it should block the activation of higher-order aggression-promoting neurons. We focused on the male-specific FruM-expressing tachykininergic neurons (Tk-GAL4^FruM^ neurons) because of their critical role in regulating overall aggression levels as well as their ability to drive robust lunging behavior when activated^14,17^. We activated Tk-GAL4^FruM^ neurons in pairs of age-matched flies using CsChrimson^19^ and then asked if the ability to induce aggression with periodic stimulation changed over time while the flies fought (Fig. 2a). Optogenetic stimulation of Tk-GAL4^FruM^ neurons initially prompted both animals to fight as expected, but as fighting continued, one animal would acutely stop responding to stimulation and switch to fleeing its opponent (Fig. 2b, Supp. Fig. 2a, Supplemental Video 2).

**Figure 2.**
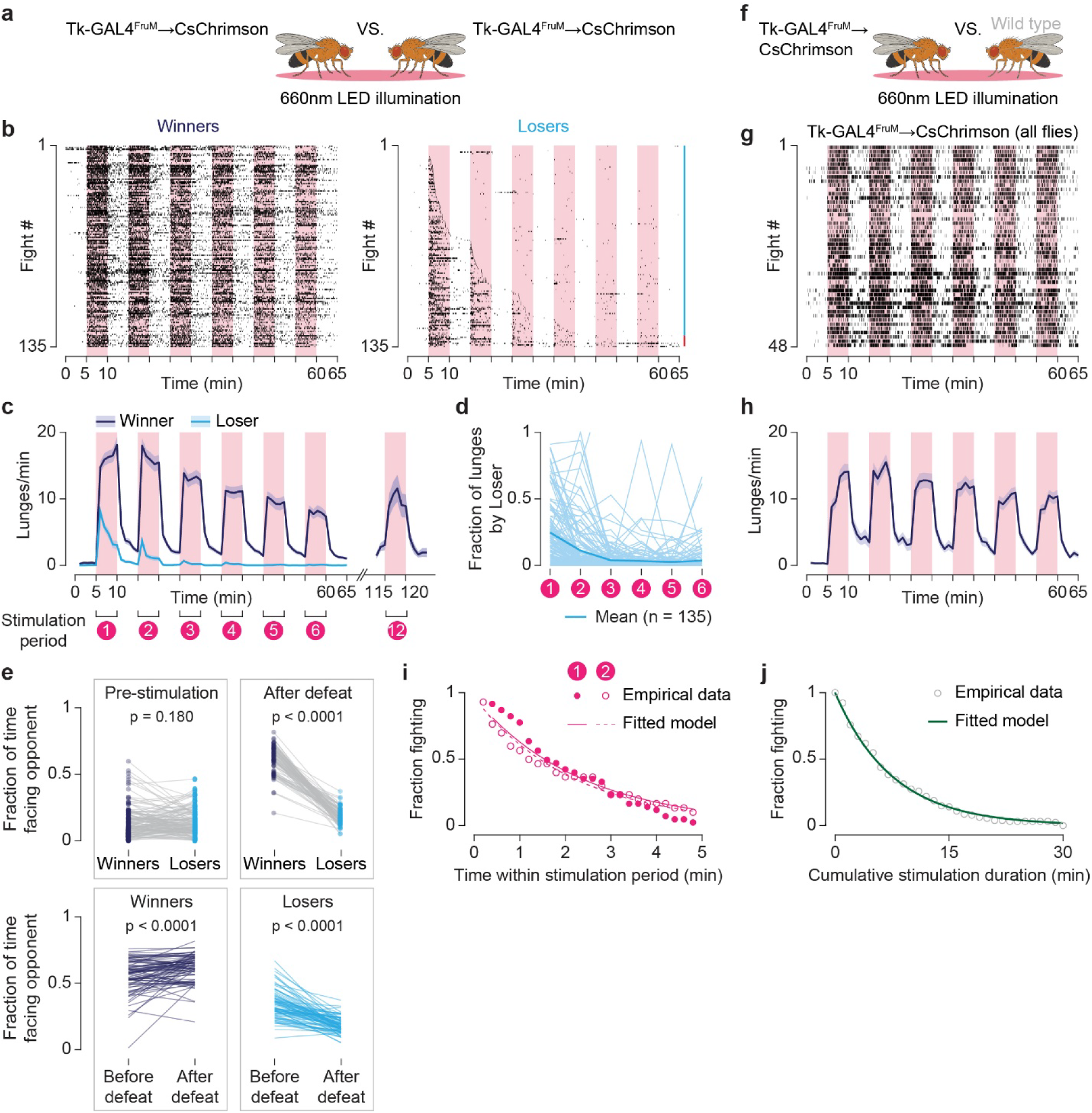
Defeat onset prevents Tk-GAL4^FruM^ neurons from eliciting aggression. **a)** Schematic of the experimental design for panels (b-d, i-j). Two age-matched males flies both expressing CsChrismon in TkGAL4^FruM^ neurons are paired to fight via optogenetic activation. **b)** Paired lunge rasters of fights induced by the optogenetic activation of Tk-GAL4^FruM^ neurons sorted by defeat time, for winners (left) and their paired losers (right). Each row corresponds to a single fly; pairs are aligned across panels by fight number. Black ticks indicate individual lunges. Pink bars indicate stimulation periods. Cyan side-bar marks loser flies that experienced defeat (see methods), red side-bar marks flies that lunged less than their opponent but didn’t exhibit defeat within the recording. n = 135 fights. **c)** Mean lunge rates of winners (dark blue line) and losers (cyan line) in (b) that were defeated by the end of the recording, n = 124 pairs, (1-min bins). Right: mean lunge rate of winners upon the 12^th^ 5-min ON/OFF stimulation period; separate dataset, n = 40 winners. Trace shading = S.E.M. **d)** The fraction of lunges by losers or low-lungers relative to total lunges per pair performed during each stimulation period. n = 135 fights (the same data set as in b). **e)** Fraction of time facing opponent (0-43° degrees). Top right: during pre-stimulation period (0-5min before first stimulation), n = 135 pairs as in (b). Top left: comparison between winners and losers after defeat. Bottom panels: comparisons among winners and losers, before and after defeat. “Before-” and “after- defeat” categories are all video frames during stimulation either before or after the defeat onset time respectively. Flies without a minimum of 1-min pre-defeat were excluded, n = 81 pairs. Median before time duration = 2.08 min, median after time duration = 6.40 min. p-values calculated by paired t-tests. **f)** Schematic of experimental design for panels (f-g): Tk-GAL4^FruM^ → CsChrimson flies paired to fight non-aggressive, group-housed Canton-S male flies. **g, h**) Lunge raster plot of Tk-GAL4^FruM^ → CsChrimson flies (g) and their mean lunge rate over time (h) (dark blue line, 1-min bins), n = 48. Trace shading = S.E.M. **i)** Fraction of fighting flies over time (12-s bins) in stimulation periods 1 (closed circles, n = 41) and 2 (open circles, n = 29), fit by an exponential function. τ = 2.35min for stimulation period 1 (R^2^ = 0.928), τ = 2.18min for stimulation period 2 (R^2^ = 0.974). **j)** Proportion of losers still fighting across all stimulation periods (circles, 1-min bins), fit by an exponential function (τ = 7.57 min, 95% CI = 7.38–7.76), n = 128 losers (excludes “instant defeat” flies that didn’t initially fight back, see Methods).

The resulting winner-loser relationship appeared similar to fights between socially isolated flies. Winners remained responsive to Tk-GAL4^FruM^ neuronal stimulation, consistently executing lunges in response to stimulation for hours, whereas Tk-GAL4^FruM^ neuronal stimulation in the eventual losers continuously failed to trigger luging following defeat (Fig. 2c, d). Following defeat, Tk-GAL4^FruM^ neuronal stimulation failed to drive losers to orient toward the opponent, and losers instead inverted their behavior, actively turning away in a manner consistent with the fleeing state we observed in socially isolated wild type males (Fig. 2e). We found that aggressive feedback between animals was required for this defeat effect. When flies expressing CsChrimson in Tk-GAL4^FruM^ neurons were paired with non-aggressive wild-type flies, optogenetic activation of the Tk-GAL4^FruM^ neurons invariably produced robust aggression for hours (Fig. 2f, g). Consistent with the WOA model regulating defeat, the winner’s lunging behavior in Tk-GAL4^FruM^ neurons-induced fights was not predictive of defeat onset (Supp. Fig. S2b-e), and losers could initially be the dominant attacker before succumbing to defeat (Fig. 2d). Retesting the same pairs of opponents 4 hours later showed that defeat largely suppressed the output of Tk-GAL4^FruM^ neurons for hours, although some losers could recover and become the dominant attacker (Supp. Fig. S2f). Furthermore, when we paired winners from the first round of Tk-GAL4^FruM^ neuronally-stimulated fights to fight each other, a new winner–loser relationship reliably emerged in the second round (Supp. Fig. 2g, h). These experiments show that both losing and winning in these contests are not a fixed trait. Together, this indicates that a winner-loser relationship arises between flies in fights driven by the activation of the Tk-GAL4^FruM^ neurons, indicating that defeat onset resulting from active fighting blocks the aggression-promoting effects of Tk-GAL4^FruM^ neuron activation.

We next measured the defeat onset times of losers in Tk-GAL4^FruM^-stimulated fights during the first 5-minute stimulation window and found that defeat onset followed an exponential distribution (Fig. 2i, solid dots). Defeat onset in the subsequent stimulation window, as well as in the cumulative stimulation duration, also displayed an exponential distribution, consistent with a model in which internal costs accumulate linearly over time (Fig. 2i, open dots, Fig. 2j). Using a continuous CsChrimson stimulation protocol instead of a pulsed protocol yielded similar results, producing an exponential defeat onset distribution with a rate-constant that approximately matched our pulsed-protocol (Supp. Fig. S2i, j). Together, our results suggest that the defeat onset distribution results from a specific circuit mechanism that probabilistically suppresses the output of Tk-GAL4^FruM^ neurons according to an exponential distribution described by evolutionary game theory.

We hypothesized that this might occur by directly suppressing Tk-GAL4^FruM^ neurons. To test this, we first generated winners and losers from optogenetically induced fights between pairs of flies that express CsChrimson and GCaMP7s^20^ in Tk-GAL4^FruM^ neurons. We then used 2-photon live imaging to measure calcium influx in the axon termini of Tk-GAL4^FruM^ neurons^21^ in response to CsChrimson-induced depolarization (Fig. 3a, b). We found that peak GCaMP7s fluorescence in losers decreased by ∼50% in response to CsChrimson stimulation compared to both winners and conflict-naïve flies without a fighting partner (Fig. 3c, d; Supp. Fig. S3a). Since CsChrimson directly depolarizes Tk-GAL4^FruM^ neurons, this reduction in GCaMP7s signal likely reflects inhibition of presynaptic calcium influx and subsequent neuronal output. Calcium influx in winners was comparable to control flies that didn’t experience fighting, suggesting that Tk-GAL4^FruM^ neurons are not modulated by the act of winning (Fig. 3c, d; Supp. Fig. S3a). These results suggest defeat onset acts in part by reducing the excitability of Tk-GAL4^FruM^ neurons, thereby decreasing the animal’s aggressive arousal to continue fighting. We hypothesized that enhancing Tk-GAL4^FruM^ neuronal activity should then prolong fighting before defeat onset. To test this, we asked whether increasing stimulation power on Tk-GAL4^FruM^ neurons could prolong fighting before defeat onset. While our standard stimulation protocol already saturated the number of evoked lunges and thus did not increase winners’ lunge rates further (Fig. 3e; Supp. Fig. S3b-d), increasing the stimulation power markedly prolonged defeat onset in losers (Fig. 3f-i). Together, our results highlight that Tk-GAL4^FruM^ neurons are subject to context-dependent modulation and support the hypothesis that the defeat and subsequent loss of aggression in losers occurs through the suppression of Tk-GAL4^FruM^ neurons.

**Figure 3.**
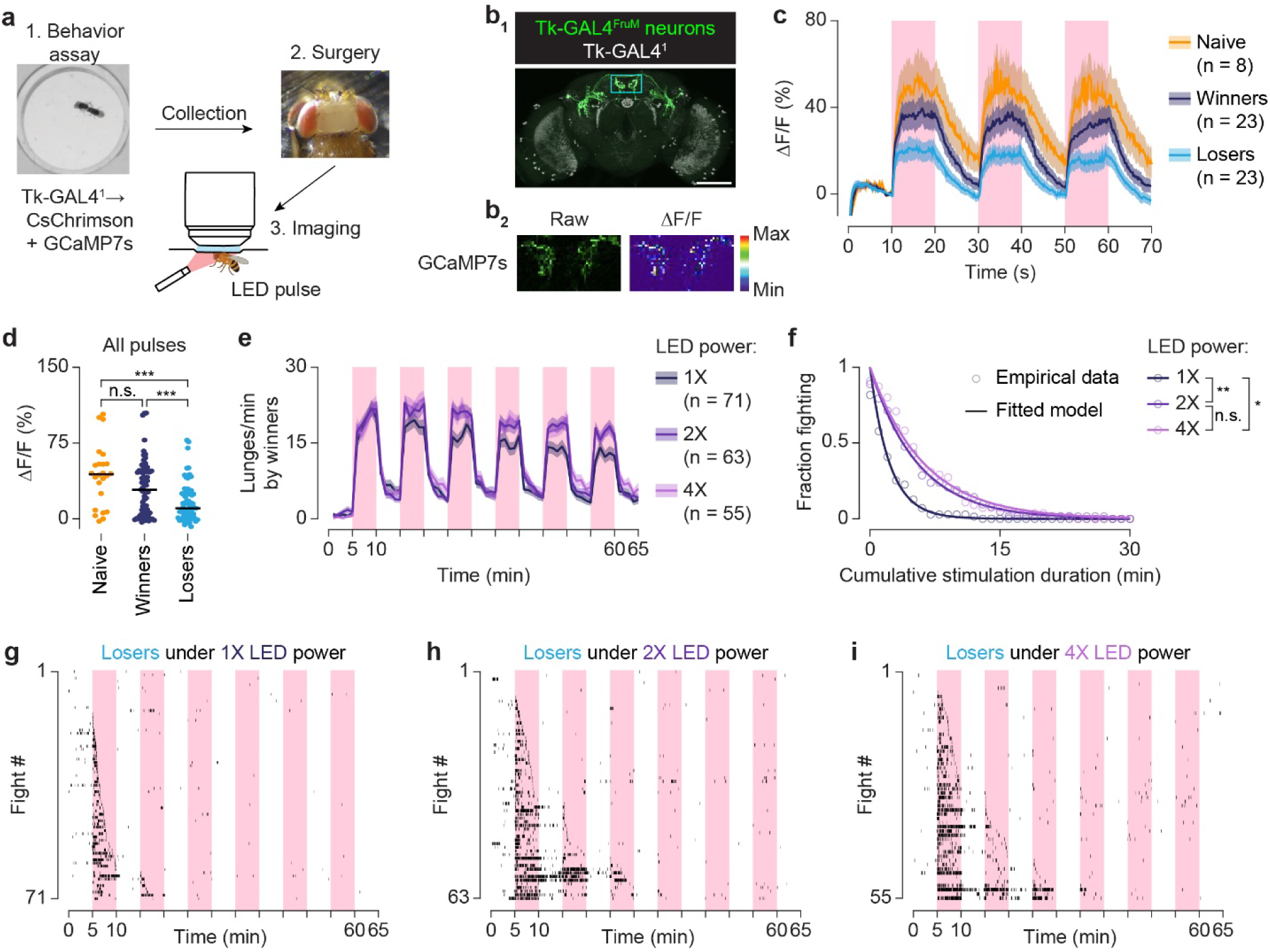
Defeat inhibits Tk-GAL4^FruM^ neuronal activity. **a)** Schematic of the calcium imaging protocol. Winners and losers were collected after fights induced by the optogenetic activation of Tk-GAL4^FruM^ neurons. Neuronal calcium influx was evoked by external 625 nm laser stimulation, calibrated to match the light intensity used during behavioral experiments. **b)** Anatomy of Tk-GAL4^FruM^ neurons and example response to optogenetic activation in a winner fly. **c)** GCaMP7s responses in Tk-GAL4^FruM^ neurons to optogenetic activation in naive (no opponent in fighting chamber, orange), winners (dark blue) and losers (cyan). **d)** Peak GCaMP7s fluorescence responses of Tk-GAL4^FruM^ neurons in each stimulation period (naïve n = 24, winners n = 69, loser n = 69). *** p < 0.001 by one-way ANOVA and *post-hoc* Tukey’s (HSD) test. **e)** Lunge rates of winners from fights induced by optogenetic activation of Tk-GAL4^FruM^ neurons using 1X, 2X, and 4X LED stimulation intensities. Increasing light intensity did not significantly increase lunge rates in all stimulation periods except for stimulus 5 and 6, 1x vs 2x LED power p < 0.05. One-way ANOVA, Tukey Correction, ns = p < 0.05. **f)** Fraction of losers still fighting over time (solid dots, 1-min bins), fit by exponential decay (solid lines). Data from (g-i), instant defeat flies excluded (1X n = 58, 2X n = 56, 3X n = 46). n.s. p = 0.76, * p = 0.01, *** p = 0.002 by pairwise Kolmogorov–Smirnov tests with Bonferroni correction. **g-i**) Lunge rasters of losers from contests in (e), sorted by time of defeat onset. Pink shaded areas indicate stimulation periods, shading around traces = S.E.M.

### A mushroom body circuit regulates defeat onset

We next sought to identify neurons important for triggering defeat in fights induced by Tk-GAL4^FruM^ activation. Silencing neurons required for defeat under these conditions should then result in contests where both flies continue fighting in response to Tk-GAL4^FruM^ activation. We used the LexA/LexAop system to orthogonally express temperature-sensitive dyamin mutant protein (Shibire^TS^)^22^, allowing us to conditionally block synaptic transmission during fighting from candidate neurons while simultaneously activating Tk-GAL4^FruM^ neurons (Fig. 4a).

**Figure 4.**
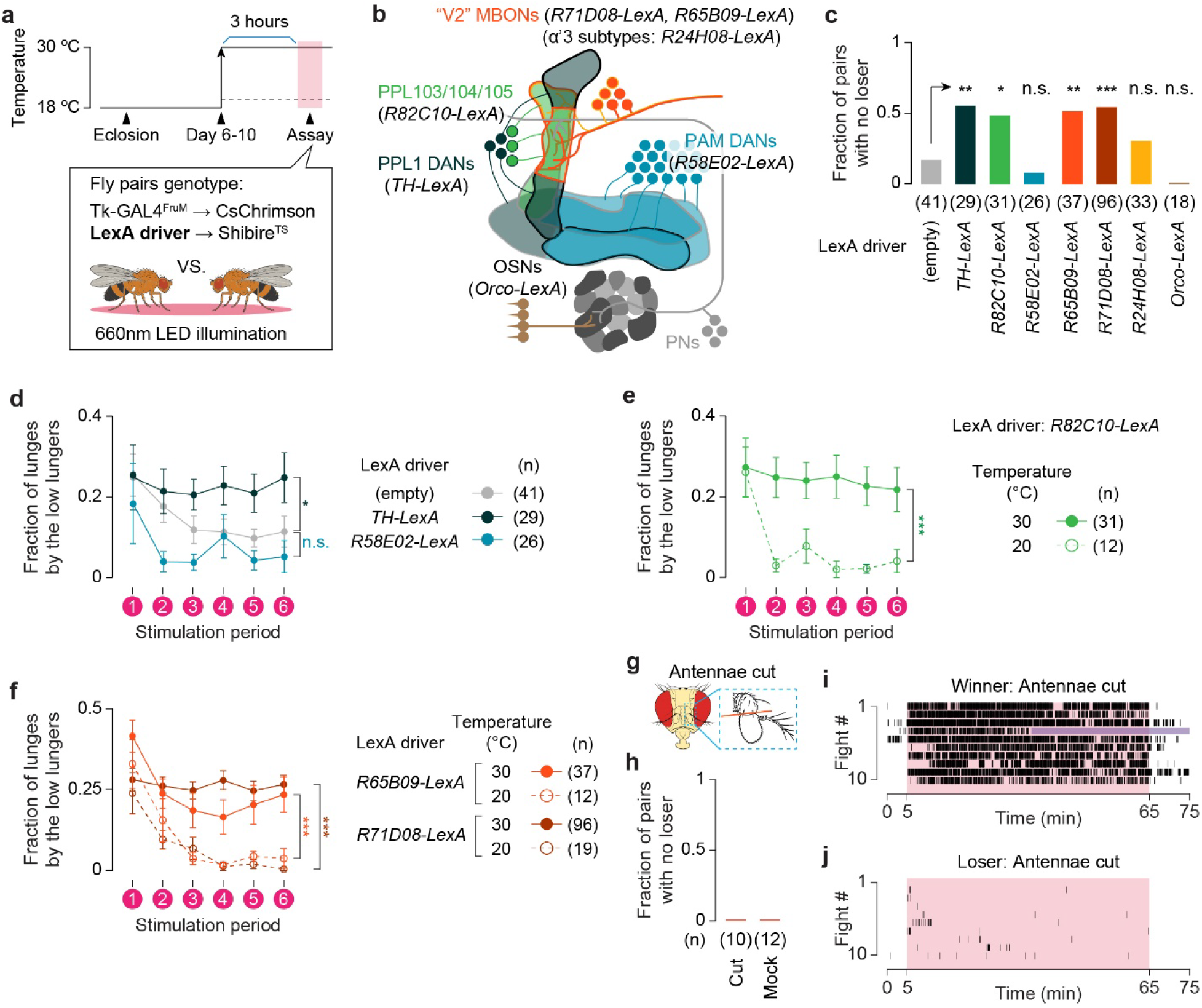
Silencing PPL1 DANs and V2 MBONs disrupts dominance formation. **a)** Diagram of fighting assay that couples optogenetic activation of Tk-GAL4^FruM^ neurons with Shibire^TS^-mediated blockade of synaptic transmission in neurons labeled by an orthogonal LexA driver. Fights were conducted for 1-hour with alternating 5-min ON/OFF light stimulation, as in previous assays. **b)** Circuit diagram of neurons labeled by the LexA lines used in these experiments. **c)** Fraction of fights that did not result in winner-loser hierarchy for each LexA driver line tested. Statistical tests are in comparison to “no LexA” genetic control (empty), Chi-square test with Bonferroni correction. **d)** Fraction of lunges performed by the fly with fewer total lunges (“low lunger”) when dopaminergic neurons were silenced. Fraction of lunges during the combined last two stimulation periods (corresponding to the dominance determination window as described in Methods) was analyzed with ANOVA and post-hoc Dunn’s test in comparison to the “no LexA” genetic control. **e, f)** Fraction of lunges performed by low lungers when PPL1 dopaminergic neurons (**e**) or V2 MBON neurons (**f**) were silenced. Welch’s t-tests between restrictive (30°C) and permissive (20°C) temperatures for each genotype of the combined last two stimulation periods. **g)** Diagram of fly head showing site of antennal removal. **h)** Fraction of fights that did not result in winner-loser hierarchy after antennal removal or mock surgery. All pairs formed dominance. **i, j**) Winner (i) and loser (j) lunge rasters from fights induced by the optogenetic activation of Tk-GAL4^FruM^ neurons between flies without antennae. Shaded pink area indicate stimulation period (1-hour). Purple shading indicates absence of lunging because the loser escaped onto the wall of fighting chamber. All panels, error bars = S.E.M., n.s. p > 0.05, * p < 0.05, ** p < 0.01, *** p < 0.001.

We first tested dopaminergic neurons given dopamine’s central role in learning, motivation, and regulation of state-dependent activity. In particular, dopaminergic neurons projecting to the mushroom bodies (DANs) from the protocerebral posterior lateral (PPL1) cluster (Fig. 4b) are important for encoding aversive signals, which we thought could be relevant for encoding cost during fighting. We found that silencing subsets of dopaminergic neurons labeled by the *TH-LexA* driver, which include the PPL1 cluster^23^, prevented formation of winner-loser relationship in 55% of contests, in contrast to only 17% among controls (Fig. 4c), resulting in contests in which both flies continued fighting in response to stimulation throughout the assay (Fig. 4d; Supp. Fig. S4a, b). Targeted silencing of PPL1 DANs using *R82C10-LexA*^24^ also disrupted defeat onset compared to permissive temperature controls (Fig. 4c, e; Supp. Fig. S4c, i; Supplemental Video 3). In contrast, silencing DANs from the protocerebral anterior medial (PAM) cluster, known for its role in encoding “reward” signals^25^, had no effect on defeat (Fig. 4c, d; Supp. Fig. S4d).

The PPL1 DANs sends axonal projections to the vertical lobes of the mushroom body (MB), where they signal to V2 mushroom body output neurons (MBONs) that mediate aversive learning in olfactory paradigms^26,27^. We found that silencing V2 MBONs using *R65B09-LexA*^28^ also blocked defeat onset in 51% of contests (Fig. 4c, f; Supp. Fig S4e; Supplementary Video 4). The V2 MBON region consists of five distinct cell-types (MBON-15 through MBON-19)^26,29^. Silencing V2 MBONs labelled by *R71D08-LexA*^28^ also blocked defeat onset (Fig. 4c, f; Supp. Fig. S4f), but silencing with *R24H08-LexA* (which labels MBON15-17)^28^ did not (Fig. 4c; Supp. Fig. S4g), suggesting that MBON-18 and MBON-19 are the relevant cell-types within the V2 cluster.

The PPL-V2 MBON circuit is known classically as being required for avoidance behavior in aversive olfactory learning paradigms. To ask if defeat onset is regulated by olfactory information, we surgically removed the antennas from flies expressing CsChrimson in Tk-GAL4^FruM^ neurons (Fig. 4g). We found that optogenetically induced fights resulted in winner-loser hierarchies similar to controls (Fig. 4h-j; Supp. Fig. S4j). Similarly, silencing *Orco*-expressing olfactory sensory neurons^30,31^ using Shibire^TS^ had no effect (Supp. Fig. S4h). These results suggest that the PPL1-V2 MBON circuit regulates defeat onset independently of olfactory input, rather than exclusively functioning in aversive olfactory conditioning. Together, these results reveal that the action to stop fighting and flee requires a mushroom body microcircuit involving specific DANs and MBONs. Without this pathway, flies continued fighting in response to activation of Tk-GAL4^FruM^ neurons, highlighting that defeat onset requires an active mechanism to block aggression output.

### Dopamine and V2 MBON activity controls defeat probability

The effect of silencing the PPL1-V2 MBON circuit can be interpreted as an increase in the perceived net payoff of fighting in the WOA model (*v* in eq. 1), resulting in flies that fight longer before defeat onset. We hypothesized that activating the PPL1-V2 MBON pathway would have the opposite effect by increasing perceived cost and biasing flies toward early defeat onset and losing. To test this, we first optogenetically co-activated either PPL1 neurons or V2 MBONs with Tk-GAL4^FruM^ neurons and found that activation of these pathways themselves did not impact aggression toward non-aggressive, group-housed wild type males (Supp. Fig. S5a-c). Optogenetic stimulation produced consistent lunging toward the wild type males, similar to animals that express CsChrimson only in Tk-GAL4^FruM^ neurons. We then paired these flies against male flies expressing CsChrimson only in Tk-GAL4^FruM^ neurons, which would fight back in response to stimulation, and asked how co-activation of PPL1 or V2 MBONs impacted the winner-loser outcomes of these competition contests (Fig. 5a). Strikingly, activation of either population significantly biased flies to lose fights (Fig. 5b, c), with V2 MBON co-activation resulting in defeat in 94% of flies (n = 69). Similar to our Shibire-mediated silencing experiments, co-activation with *R24H08-LexA* had no effect on winner-loser outcomes (Fig. 5d; Supp. Fig. S5d).

**Figure 5.**
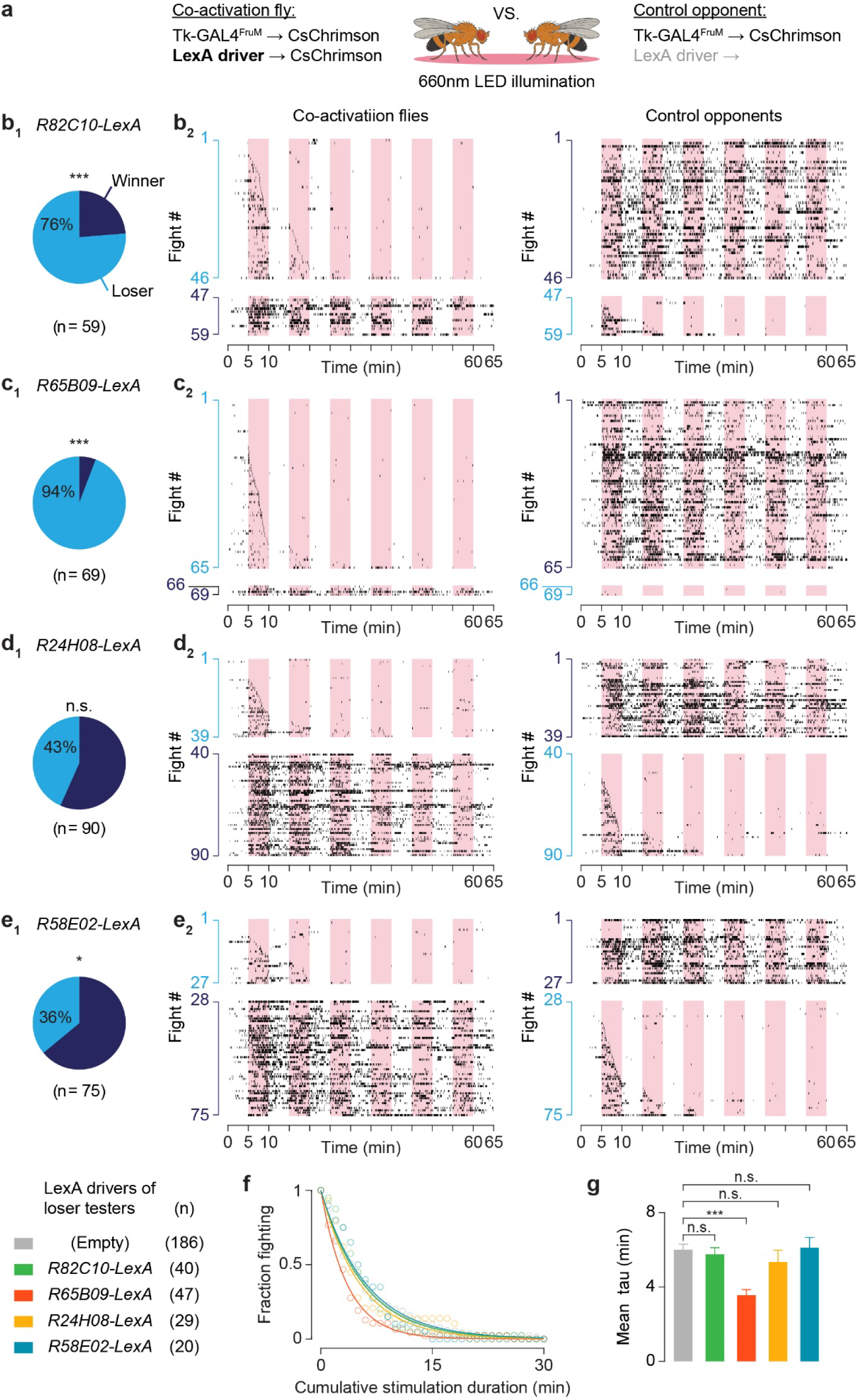
Activation of dopaminergic and MBONs reshapes dominance formation. **a)** Schematic of competition assays designed to test how activating dopaminergic and MBONs affects winner–loser outcomes in fights induced by the optogenetic activation of Tk-GAL4^FruM^ neurons. In these assays, flies in which LexA-labeled neurons were co-activated with Tk-GAL4^FruM^ neurons were paired against control opponents in which only Tk-GAL4^FruM^ neurons were stimulated. **b-e**) Outcomes of fights between co-activation flies and control opponents. (**b_1_-e_1_**) Breakdown of co-activation flies that became winners and losers. Binominal test. n = number of fights tested. (**b_2_-e_2_**) Paired lunge rasters for co-activation (left) verse control opponents (right) separated by winner-loser outcomes, indicated by colored y-axes (cyan = losers, dark blue = winners). Rasters are sorted by defeat onset time. Pink bars in raster indicate CsChrimson stimulation periods. **f)** Proportion of losers still fighting over time (open circles, 1-min bins), fit by exponential decay (solid lines). **g)** Mean time constants (τ) from fits to bootstrapped distributions of defeat onset times as in (f). Error bars are S.D. Welch’s unpaired t-tests with Bonferroni correction. Stars mark significant comparisons; all unmarked comparisons are not significant. Sample number (n) reflects loser flies used for fitting (see Methods). All panels: n.s. p < 0.05, ** p < 0.01, *** p < 0.001.

According to the WOA model, an elevated internal cost should reduce the time constant of the exponential distribution governing defeat onset, resulting in a faster decay of the probability distribution. Consistent with this prediction, V2 MBON co-activation resulted in defeat onset distributions that were markedly left-shifted compared to losers that only express CsChrimson in Tk-GAL4^FruM^ neurons (Fig. 5f, g). PPL1 co-activation did not strongly affect the defeat onset distribution, suggesting that PPL1 neurons may play a more modulatory role in the circuit or impact contest outcomes through additional mechanisms.

Since aversive signaling pathways appeared to be important for regulating defeat, we hypothesized that the activation of the PAM DAN reward pathway might instead increase fighting persistence to facilitate winning. Similar to the PPL1-V2 MBON circuit, co-activation of PAM neurons did not impact aggression induced by the optogenetic activation of Tk-GAL4^FruM^ neurons when these flies were paired with non-aggressive wild-type opponents (Supp. Fig. S5e). However, this manipulation did bias these flies toward winning in optogentically induced fights against opponents that expressed CsChrimson only in Tk-GAL4^FruM^ neurons (64% won, n = 75; Fig. 5e). Together, these results indicate that dopamine signaling from distinct pathways can play a dual role in controlling contest outcomes.

### Internal state modulates defeat onset probability

Under the WOA model, payoff is represented by the decay constant of the exponential probability distribution. Individuals are more likely to flee from the fight if the cost of persisting in the contest is high relative to the resource’s value. This would be represented by a steeper exponential decline in the probability function. Conversely, if the resource is highly valuable, individuals adopt overall longer waiting times, represented by a flatter exponential distribution function. In *Drosophila*, food is a resource commonly required to induce fighting^6,32–34^. This suggests food presence increases perceived payoff of initiating social conflict. Yeast is an important food resource for flies that is also known to increase aggression^6,34^. Flies deprived of protein in their diet show a strong preference for yeast^35,36^, suggesting that they assign a higher value to yeast than fully fed flies. This suggests that hunger state can change the perceived value of a food resource as well.

To ask if perceived payoff impacts defeat onset accordingly, we investigated how hunger state and the presence of a food resource impacts the probability distribution of defeat onset times. To do this, we measured the defeat onset distributions in fights driven by the activation of the Tk-GAL4^FruM^ neurons under fed and starved conditions, either with or without the presence of yeast (Fig. 6a). Consistent with the WOA model, defeat onset distributions shifted systematically across hunger and food resource conditions and were well described by changes in the decay constant (*v* in eq.1) of an exponential fit (Fig. 6b-g). These changes in the distributions also aligned with behavioral predictions based on the effects of perceived payoff. Measuring the EC_50_ values from our exponential fits, we found that starved flies fought longer than fed flies, and starved files in the presence of yeast persisted longer than starved flies without yeast (Fig. 6c). These results support that internal state modulates contest persistence by tuning the probability of defeat, consistent with the role of modulating payoff in the WOA model.

**Figure 6.**
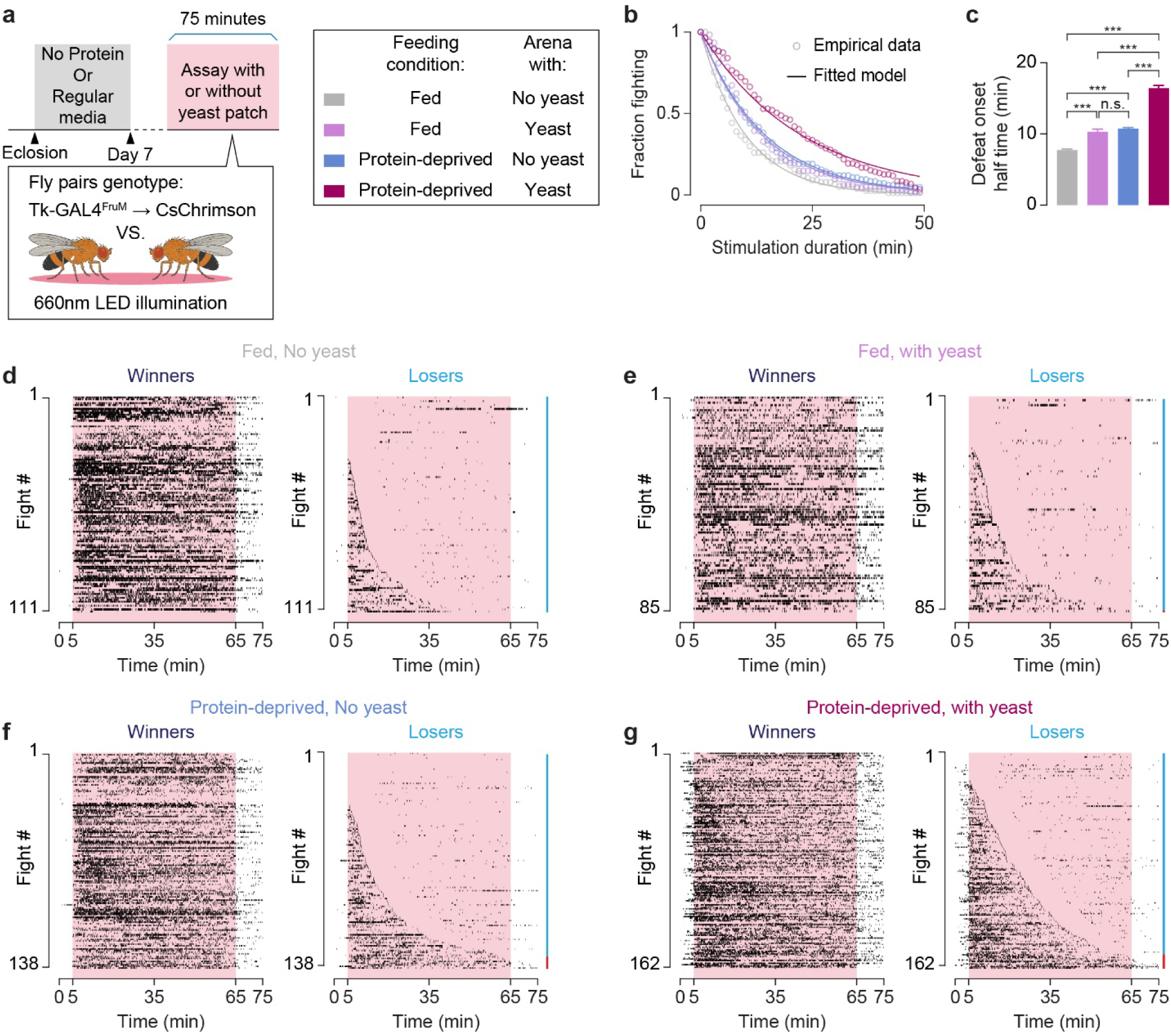
Internal state and resource-presence modulate defeat onset probability. **a)** Left: Schematic diagram of fighting assays designed to test how protein deprivation and/or resource presence influence fighting induced by Tk-GAL4^FruM^ stimulation. After eclosion, CsChrimson-expressing flies were reared on either standard or sucrose media (no protein) for 7-days and then paired to fight in chambers with or without a patch of yeast. Right: Legend diagram of conditions tested. **b)** Proportion of losers still fighting over time for each condition (open circles, 1-min bins), fit by exponential decay (solid lines). **c)** Mean half-life (*t*_1/2_) from fits to bootstrapped distributions of defeat onset times as in (b). Error bars are S.D. Welch’s unpaired t-tests with Bonferroni correction. n.s. p < 0.05, *** p < 0.001. In (b) and(c), pairs with instant defeats or no dominance are removed (see Methods): Fed without yeast, n = 77; Fed + yeast, n = 61; Protein deprived without yeast, n = 84; Protein deprived + yeast, n = 119. **d-g**) Paired winner-loser lunge rasters for each condition tested. Cyan side-bar marks loser flies that experienced defeat, red side-bar marks flies that lunged less than their opponent but didn’t exhibit defeat within the recording. Pink bars in raster indicate CsChrimson stimulation periods (1hr of 10ms pulses at 5Hz).

## DISCUSSION

The evolutionary logic of competitive aggressive behavior predicts that animals should adopt strategies that balance the benefits of winning with the costs of fighting. Our findings provide evidence that such principles are embedded within the neural circuitry that regulates aggression, providing a mechanistic link between evolutionary theory and neural functions.

Dominance formation in *Drosophila* has been noted since the earliest descriptions of fly aggression^37,38^, but the neural processes that regulate winner–loser outcomes from fighting has remained unclear. The choice to cease fighting and flee in male *Drosophila* is the key behavioral transition that determines dominance outcomes^8,11^. We find that this choice follows a strategy that is an evolutionarily stable solution to War-of-Attrition type contests. A basic assumption of this model is that the choice to flee is regulated through a probabilistic selection of a persistence-time sampled from an exponential distribution, rather than being driven directly by the opponent’s actions. Our results are consistent with this model and with previous work showing that aggressive actions including lunging do not deterministically trigger dominance formation^11^. Instead, we propose that contest outcomes are determined by the interaction of underlying persistence-time probability distributions sampled by each opponent, which themselves are shaped by the individual’s internal state (e.g. hunger-state). This provides a mechanistic explanation for both why asymmetries in contest outcomes can arise and why they remain inherently variable.

Our circuit-level dissection identified the mushroom body as a core neural component underlying this process. We hypothesize that mushroom body circuitry regulates defeat onset by integrating internal states and generating a probabilistic output resulting in the observed exponential distribution of defeat times, in accordance with the WOA model described by game-theory (eq. 1). This interpretation is broadly consistent with a growing body of work showing that the mushroom body organizes probabilistic behaviors, including social behaviors, by integrating competing sensory inputs, internal states, and reinforcement signals^39–41^. Within this framework, payoff in the WOA model may be represented within mushroom body circuits as the integrated value of internal states, consistent with our finding that hunger and resource availability modulate the probability of defeat onset.

Locomotion and movement have also been shown to be represented in the activity states of both the mushroom body and dopaminergic neurons, suggesting that these circuits can integrate ongoing motor behavior into decision-making and goal-directed processes^42–45^. This observation is relevant to our finding that the PPL1–V2 MBON circuit seems to be recruited to suppress Tk-GAL4^FruM^ neurons’ function only after receiving agonistic signals from active fighting. PPL1 neurons are activated by negative stimuli such as electric shock^46^, and dopamine released from these neurons is essential for aversive learning through modulation of MBON activity^47,48^. This suggests that PPL1 activity during fighting could represent a negative valence, shaping decision-making in the context of social conflict. We hypothesize that PPL1 signaling in response to agonistic cues modulates V2 MBON activity, thereby triggering avoidance behavior^26,27,49^.

Defeat onset suppresses aggression resulting in flies that actively flee their opponent. We identify Tk-GAL4^FruM^ neurons as an important node in this process in male flies. These neurons, previously implicated in promoting aggressive arousal^14^, appear to function as a gate point for defeat onset via their inhibition. This inhibition depends on PPL1 DANs and V2 MBONs; however, inhibition is likely indirect. Both PPL1 neurons and V2 MBONs do not make direct connections with Tk-GAL4^FruM^ neurons (Supp. Fig. S6a-c), and their activation alone does not inhibit Tk-GAL4^FruM^ neurons from inducing aggression. This is consistent with previous work showing that PPL1 neurons do not impact spontaneous male aggression, but dopamine from other sources modulates aggression arousal levels^50^. This again emphasizes that the PPL1-V2 MBON circuit’s interaction with aggression circuits is context-dependent and is likely recruited through aggressive feedback.

Following defeat, Tk-GAL4^FruM^ neurons enter a prolonged suppressed state that can persist for hours, during which optogenetic activation no longer triggers aggression. This aligns with previous work showing that defeat can induce long-lasting reductions in male aggression in *Drosophila*^8–11^, and suggests that such effects result from sustained suppression of Tk-GAL4^FruM^ neurons. These long-lasting effects are also reminiscent of the enduring behavioral effects seen after repeated social defeat stress in both insects and mammals^51–54^. The mechanism that triggers the choice to stop fighting can be distinct from the mechanisms underlying this type of long-lasting inhibition, although in male flies both may converge on the suppression of Tk-GAL4^FruM^ neurons at different time scales. In our current experimental paradigm, the loser flies cannot escape from the chamber, potentially resulting in a condition similar to “repeated social defeat”. Neural mechanisms underlying the transition from the defeat onset to a longer-lasting state change is an interesting topic for future study.

Our work provides a new framework for interpreting how neural circuits resolve conflicts across species. In zebrafish, dorsal habenula (dHb) circuits influence contest outcomes independently of aggression: individuals that ultimately lose can initially be the dominant attacker before switching to defeat^55^. Starvation further modulates this pathway, increasing fighting duration and persistence and thereby enhancing the likelihood of victory^56^. These parallels suggest that dHb circuitry may implement a computation analogous to what we describe in this study. In rodents, social dominance is orchestrated by a distributed network of brain regions^57,58^. Yet, the mechanisms that govern the decision to halt fighting and withdraw, especially when opponents are evenly matched, remain unresolved. Recent work highlights prefrontal–thalamic circuits as critical regulators of dominance formation, demonstrating how top-down control integrates internal state and social context to shape competitive behavior^59–62^. Our findings raise a possibility that probabilistic computations may be embedded within these networks to implement cost–benefit decisions that support evolutionarily stable strategies^63^.

Our work provides mechanistic evidence that natural selection has embedded game-theoretic principles into the neural substrates governing social conflict. By linking evolutionary models of competition with the cellular and circuit-level mechanisms that drive behavior, we reveal how fundamental adaptive strategies can be implemented in the brain, offering a framework for understanding the neural basis of a strategic action choice during social interactions.

## Supporting information

Raw Data file for main figures

Raw Data file for supplemental figures

## ACKNOWLEDGEMENTS

We thank Juliet Heller, Henry Kwan, and Ezgi Yalbir for their assistance on behavioral and immunohistological experiments, and Steven Flavell and Daisuke Hattori for critically commenting on the manuscript. This work has been supported by NIH NIGMS R35GM119844 to K.A. D.V. was supported by a Pioneer Fund Postdoctoral Scholar Award of the Salk Institute, and by a Kavli Institute for Brain and Mind Postdoctoral Award.

## AUTHOR CONTRIBUTIONS

D.V. conceptualization, methodology, software, investigation, validation, formal analysis, data curation, writing, visualization, funding acquisition; E.C.-C. Investigation; C.S.L. Investigation; B.R. methodology, software, investigation; K.A. conceptualization, investigation, validation, formal analysis, data curation, writing, visualization, supervision, funding acquisition.

## MATERIALS AND CORRESPONDENCE

Requests for materials should be directed to K.A.

## MATERIALS AND METHODS

### Experimental Animals

Canton-S, *Tk-GAL4^1^* (ref. #14) (RRID:BDSC_52740), *Otd-nls:FLPo* in attP40, *20XUAS>myr:TopHAT2>Syn21-CsChrimson:tdTomato* in attP2 (ref. #14), *fru^FLP^* (RRID:BDSC_66870) (ref. #64), and *20xUAS*>*myr:TopHAT2*>*GCaMP6f* in su(Hw)attP5 (ref. #21) were gifts from David Anderson (Caltech). *13xLexAop2-IVS-Syn21-shibire^TS^* in VK00005 (ref. #65) was a gift from Gerald Rubin (HHMI Janelia Research Campus). *TH-LexA*^23^ was a gift from Kristin Scott (UC Berkeley). *20X-UAS-IVS-jGCaMP7s* in su(Hw)attP5 (ref: #20) (RRID:BDSC_80905), *R58E02-LexA* in attP40 (ref: #25) (RRID:BDSC_52740), *R82C10-LexA* in attP40 (RRID:BDSC_54981), *R65B09-LexA* in attP40 (RRID:BDSC_52687), *R71D08-LexA* in attP40 (RRID:BDSC_52841) (ref. #28), *R24H08-LexA* in attP40 (ref. #24) (RRID:BDSC_52732) were obtained from Bloomington Drosophila Stock Center (Indiana University). *Orco-LexA*^66^ was a gift from Jing Wang (UC San Diego). See Supplementary Table for the complete genotypes of transgenic flies used in this study.

### Animal Preparation

Experimental flies were grown on standard cornmeal-based food at 25°C with 60% relative humidity (RH) on a 12hr light/dark cycle. Flies were collected using CO_2_ anesthesia on day of eclosion. CO_2_ exposure was strictly limited to less than 4min. During collection the tips of a wing were clipped using a razor blade to facilitate individual identification during behavior experiments. Experiments were performed between 6-10 days from eclosion. Flies were transferred to fresh food vials every 3 days. For optogenetic experiments, flies were transferred to food supplemented with 200μM all-*trans* retinal (20 mM stock solution prepared in 95% ethanol; cat# R2500, Millipore Sigma) 2–3 days before experiments. These flies were shielded from light until the day of experiment. Group-housed flies contained 8-18 flies per vial. Flies used in Shibire^TS^ experiments were kept at 18°C and were shifted to 30°C or 20°C for ∼3 hours prior to behavior experiments. Most behavior assays were conducted during Zeitgerber time (ZT) 7-12, with some experiments done during ZT 0-3.

For yeast-deprivation experiments, on the day of eclosion flies were transferred to vials containing agar and 200mM sucrose. Flies were transferred to fresh sucrose vials every 3 days until behavioral experiments, which were conducted 7-10 days after eclosion.

For antennae removal experiments, 5-day old adult male flies were anesthetized on CO_2_ for 30-seconds during which their antennae were removed using sharp microforceps. Mock-treated controls underwent the same anesthesia without antennae removal. Flies then were allowed to recover for 2-days on media containing 200μM of all-*trans* retinal before testing.

### Behavioral Assays

Behavior assays were conducted at 25°C with 60% RH in climate-controlled booths except Shibire^TS^ experiments, which were conducted in either 20°C (permissive temperature) or 30-32°C (restrictive temperature) with 60% RH. Experiments were conducted in 12-well acrylic chambers^14^ using food substrate base containing 2.25% w/v agarose and 2.5% w/v sucrose in apple juice (Minute Maid) unless otherwise specified. For yeast-deprivation experiments (Fig. 6a), the substrate base contained 2.25% w/v agarose, and a 2μl drop of 5% yeast was applied to the center of each well and allowed to dry before the assay. Flies were transferred to behavior devices using aspiration directly from collection vials. To prevent flies from climbing, the chamber walls were coated with Insect-a-Slip (cat# 2871C, BioQuip Products), and the acrylic ceiling was coated with SurfaSil Siliconizing Fluid (cat# TS-42800, Thermo Fisher Scientific)^14,67^.

Behavior videos were recorded at 60fps using Flea3 USB 3.0 cameras (cat# FL3-U3-13Y3M-C, Teledyne FLIR) using a 35mm lens (cat# HF35HA1B, Fujinon) and infrared long-pass filter (cat# LP780-25.5, Midwest Optical Systems). Video images of the flies were acquired using the infrared (850nm) backlight illumination provided by custom-designed multicolor LED backlights (Metaphase Technologies Inc). In experiments that did not use optogenetics stimulation, white light was used instead of the infrared light. Behavior videos were acquired using Streampix software (version 8.5.0.0, Norpix) in MP4 format using H.264 GPU-accelerated real-time compression. Flies were discarded after each experiment and the food substrates were exchanged with fresh ones after each experiment.

### Optogenetics

To activate neurons that transgenically express CsChrimson, 660nm red-light was delivered in 10ms pulses at 5Hz at an intensity of 12μM/cm^2^ using the above-mentioned multicolor LED backlights unless otherwise stated. Power measurements were made with a photodiode power sensor (cat# S130C, Thorlabs) coupled to a digital optical power/energy meter (cat# PM100D, Thorlabs) at the surface of the behavior chamber. Optogenetic stimulation periods were custom-scripted and controlled from Streampix for acquisition synchronized stimulation: ON/OFF TTL output from Streampix were sent to an Auduino Uno R3 microcontroller board via a USB-6001 DAC (part #782605-01, National Instruments), which then outputs 10ms ON/OFF TTL signals at the specified rate (i.e. 5Hz) to the LED controller (Metaphase Tech ULC-2).

For 2-rounds optogenetically induced fighting assay (Supp. Fig. S2g), winners were collected immediately after the Round 1 via brief CO_2_ anesthetization and paired to fight against each other in Round 2 after a recovery period of 10 min.

### Quantification of behavioral data

Behavior movies were analyzed as previously described^17,68,69^. Briefly, the movies were processed by FlyTracker^70^ (version 1.0.5; https://github.com/kristinbranson/FlyTracker) and JAABA classifiers were used to detect lunges^68,71^ (https://sourceforge.net/projects/jaaba/files/). Other measurements such as facing angles and speed were calculated directly from .trx and .feat files generated by FlyTracker. Identity switch of each fly was manually corrected using the Visualizer component of FlyTracker. All behavior analysis was done in MATLAB 2024b.

Winner-loser dominance formation was identified using a three-stage analysis pipeline consisting of lunge-based classification, facing-angle classification, and a final cross-metric integration step designed to capture when a fly stopped attacking and switched into a fleeing state (see Supplemental Methods Fig 1). This analysis was performed identically across all experiments, using a 10-min dominance determination window adjusted to match the stimulation and/or recoding paradigm to classify dominance status. For pulsed CsChrimson optogenetic experiments, dominance was evaluated during the final two 5-min stimulation periods. Continuous CsChrimson stimulation experiments used the last 10-min of the stimulation window. For recordings without stimulation, we analyzed the final 10-min of the recording.

In the first stage, we evaluated dominance from per-fly facing-angle distributions. Facing angle was computed as the angle between the fly’s forward orientation vector and the vector pointing from the fly to its opponent. We focused on time points when the distance to the arena wall exceeded 3 mm to exclude constrained behaviors near the edge. Facing angles were binned from 0 to π radians and analyzed across three regions: 0–0.75 rad (*facing*), 2.4–3.14 rad (*avoiding*), and 1.2–1.8 rad (*center*). Mean differences between regions were used to determine the prevailing facing angle. Dominance formation was then marked by comparing each pair’s facing angle distributions: If both flies’ distributions peaked in the facing range, the pair was classified as ‘No Dominance’. If one fly peaked in the facing region, while the other peaked in the fleeing region, they were labelled dominant and submissive respectively. Cases with distributions dominated by the center range or lacked clear peaks were classified as ‘Unclear’. No cases were found in which both flies had peaks in the fleeing region.

In the second stage, lunge-based dominance within a pair was determined by quantifying lunges within the dominance determination window. Each pair’s high- and low-lunging individuals were identified based on their total lunge counts. If both flies failed to lunge, the pair was classified as ‘No Fighting’. If the high-lunger produced fewer than 10 lunges, the pair was designated as ‘Low Lunging’. Otherwise, dominance was assigned if the low-lunger produced ≤ 5 lunges (loser) and the total lunges of high lunger (winner) was at least twice as high as that of the low-lunger. Pairs in which each fly lunged more than 5 times were classified as ‘No Dominance’. This stage provided a robust behavioral measure of asymmetry based solely on aggressive output.

Finally, lunge- and facing-angle classifications were integrated to generate a consensus. Agreement between metrics was accepted directly. In cases where the classifications were not in agreement, decision rules prioritized the assignment of ‘Dominance’ or ‘No Dominance’ classification from either method. These calls overrode ‘Low Lunging’ classification with lunge-based method and ‘Unclear’ classification with the facing angle-based method. These circumstances were mostly relevant to cases when flies were able to climb onto the walls of the assay chamber, decreasing counted behaviors, or in Shibire^TS^ experiments, which sometimes exhibited a lower level of lunging near the end of the assay at 30°C. The output of the resulting classifications Dominance (winner/loser) and No Dominance was then used to select contests for analysis. “No Fighting” or “Unclear” pairs were not included for further analysis. For fights among socially isolated flies, pairs in which neither fly showed sustained aggression during the first 15 min (defined as fewer than five one-minute bins containing at least one lunge) were excluded (See Supp. Fig. S1a).

Defeat onset was defined as the first window in which the loser entered a defeated state (≤ 5 lunges and fleeing) for a minimum of 10-min. Winners had to be actively lunging as defined by the lunge classifier above. Defeat onset was then defined as the time of the loser’s last lunge. In most cases, this defeated state lasted for the remainder of the assay (see Fig. 1e).

### Functional imaging

Winners and losers were generated in flies co-expressing CsChrimson and GCaMP7s in Tk-GAL4^FruM^ neurons through optogenetic experiments as described in Fig. 2a, using the 5min ON/OFF stimulation protocol (see also text). Naïve control flies underwent the same protocol but lacked an opponent in the fighting chamber. At the end of the behavior assay, a fly from the winner-loser pair was collected via aspiration from the behavior chamber, briefly anesthetized on ice (∼30s), and then mounted on a custom head-fixing device using ultraviolet curing adhesive (Norland Optical Adhesive 63). The head, proboscis, thorax, and legs were secured with adhesive to prevent movement during imaging. A window for imaging was made by removing cuticle, underlying fat tissue and air sacs above the brain with forceps. Surgery and subsequent imaging were performed in *Drosophila* adult hemolymph-like saline^72^ (2mM CaCl_2_) at room temperature (∼20°C). Collection and mounting were completed within ∼5min and once mounted on the microscope, a 5 min rest period was provided before imaging. This process was then repeated for the next fly in the pair. Typically, both flies in a winner-loser pair were imaged but in some cases one fly was lost during collection or surgery. Losers of the fighting pair were typically imaged first based on brief manual inspection of the behavioral recording. Winner-loser classification was then confirmed after imaging experiment and analysis were completed. Data acquisition was limited to 1hr post behavior experiments.

Imaging was conducted on an Olympus FV-MPE-RS multiphoton laser scanning microscope, equipped with Olympus 25x water immersion objective (cat# XLPLN25XWMP2). Optogenetic stimulation was delivered by a 625 nm external fiber-coupled LED (cat# M625F2, Thorlabs). Arduino Uno Rev3 microcontroller board was used to control the LED driver (cat# DC2200, Thorlabs) and synchronize stimulation with image acquisition via TTL signals. The end of the LED fiber (cat# M28L01, Thorlabs) was placed within 1mm of the fly’s head. Stimulation power (12mW/cm^2^) was matched to behavior experiments and was supplied in 10ms pulses at 5Hz.

GCaMP7s fluorescence was imaged using a tunable laser set at 920 nm (Spectra-Physics InSight DL Dual-OL, Newport). Following the calcium imaging protocol, fluorescence of GCaMP7s and tdTomato (tagged to CsChrimson) were imaged simultaneously to aid ROI selection. CsChrimson:tdTomato was imaged with a 1040nm fixed auxiliary laser. Images were taken at ∼5Hz, depending on the size of scanning area, with a 640 × 640 pixel resolution.

Image sequences in .OIR format were converted to .TIFF and then analyzed in ImageJ (Fiji)^73^ via the Olympus ImageJ plug-in (http://imagej.net/OlympusImageJPlugin). Calcium imaging responses were extracted from the axon termini of Tk-GAL4^FruM^ neurons in the superior medial protocerebrum^21^. A custom imageJ script facilitated manual placement of ROIs to extract mean pixel intensity measurements from neuronal projections and a neighboring background region. External optogenetic stimulation resulted in red-light bleed-through into acquired images. If this coincided during scanning within a ROI, contaminated pixels (identified by scanline saturation) were removed prior to ROI measurements. GCaMP7s ΔF/F was calculated in MATLAB. Baseline fluorescence (*F*_base_) was calculated by averaging the fluorescence for 2s preceding the stimulation. ΔF/F for the frame *i* was calculated as follows:

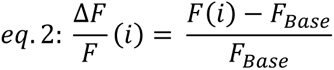

The ΔF/F for frames during the three 10s LED stimulation windows was averaged to calculate the ΔF/F mean response for each animal. Animals with excessive movement were discarded.

### Immunohistochemistry

The following antibodies, along with the indicated dilution ratios, were used for immunohistochemistry: rabbit anti-DsRed (1:1000; cat# 632496, Takara Bio; RRID:AB_10013483), chicken anti-GFP (1:1000; cat# ab13970, Abcam; RRID:AB_300798), mouse anti-Bruchpilot (1:10; cat# nc82, Developmental Studies Hybridoma Bank; RRID: AB_2392664), goat anti-chicken Alexa 488 (1:100; cat# A11039, Thermo Fisher Scientific; RRID:AB_2534096), goat anti-rat Alexa 488 (1:100, cat# A11006, Thermo Fisher Scientific; RRID:AB_2534074), and goat anti-mouse Alexa 633 (1:100; cat# A21052, Thermo Fisher Scientific; RRID:AB_2535719). Immunohistochemistry followed as described in^17,69^. Z-stack images were acquired at the Salk Biophotonics Core using a Zeiss 880 confocal microscopy, and visualized in ImageJ.

### Connectome data analysis

The male CNS connectivity data (“connectome-weights-male-cns-v0.9-minconf-0.5.feather”) was downloaded from the Male CNS Connectome Project Page^74^ (https://male-cns.janelia.org/). From this dataset, the shortest neuronal paths from 4 types of V2 MBONs (MBON17, MBON17-like, MBON18, and MBON19) and AVLP727m, a cell type that is annotated as Tk-GAL4^FruM^ neuron in MCSN v0.9, were manually identified. The minimum of 5 synapses was required to be classified as a neuronal connection. Under this threshold, none of the 1st order or the 2nd order downstream of the V2 MBONs is AVLP727m. The total number of synapses in a 4-neurons path between these 2 classes of neurons was counted, and the top 2 paths from each V2 MBON subtypes were used as representatives.

### Statistical analysis

Statistical analyses were performed in either MATLAB 2024b (Statistics and Machine Learning toolbox) or GraphPad Prism 10.4.1. Sample size was not predetermined through statistical methods before initiating the study. See the raw data file for details of statistical test results. Types of statistical tests used in figures are also indicated in figure legends. Unless otherwise indicated, statistical tests were two-tailed.

Fitting of defeat onset data was restricted to losers that exhibited fighting before defeat and met the 10-min defeated state duration stated above. Defeat onset times were used to construct a survival curve 𝑆(𝑡), defined as the fraction of individuals with defeat times greater than 𝑡, binned at 1-minute intervals. For optogenetic stimulation data, defeat onset was re-sorted using 5-min defeat-state minimum to better estimate the rate defeat onset. A single-parameter exponential decay model 𝑆(𝑡) = 𝑒^−𝑡/𝑣^ was then fitted to the survival data using nonlinear least-squares regression using MATLAB’s (2024b) in-built Fit function.

To compare decay constants and EC₅₀ values of defeat onset derived from exponential model fits, we used a subsampling bootstrapping approach to enable comparison across conditions. In each of *n* bootstrap iterations, a random fraction (75%) of defeat times was selected and re-fit to the exponential model 𝑆(𝑡) = 𝑒^−𝑡/𝜏^. This yielded distributions of τ and corresponding half-lives (𝑡𝑡_1/2_ = 𝜏ln (2)). Mean values, standard errors, and 95% confidence intervals were then derived from these bootstrap distributions. Distributions were generated using *n* = 10,000 bootstrapping iterations.

## Data availability

The raw data file contains data necessary to reproduce all figure plots. Primary data, such as videos and frame-by-frame annotations of behavioral bouts, is available upon request to corresponding authors.

## Code availability

All analysis of behavioral and imaging data was done in MATLAB 2024b. Some parts of the analysis scripts were generated with the aid of ChatGPT. These portions of code were inspected and confirmed by the first author. All code is available at: https://github.com/asahinak/Ventimiglia_etal_2025_DominanceAnalysis

Supplemental Video 1: Formation of winner-loser hierarchy during spontaneous fights between wild type male flies, related to Figure 1.

2 examples of defeat onset in fighting pairs of wild-type flies, highlighting periods before, during, and after the event of defeat onset. The fight numbers correspond to Fig. 1a and 1b. Canton-S male flies were socially isolated for 6-8 days before the assay.

Supplemental Video 2: Formation of winner-loser hierarchy during optogenetically induced fights, related to Figure 2.

2 examples of defeat onset in fighting pairs of Tk-GAL4^FruM^ → CsChrimson flies, highlighting periods before, during, and after the event of defeat onset. The fight numbers correspond to Fig. 1b and Supplemental Fig. S2a. Note that in fight #70 (bottom), the eventual loser fly lunged more than the eventual winner during the active fighting.

Supplemental Video 3: Blocking the synaptic transmission of PPL1 DA neurons disrupts defeat onset, related to Figure 4.

Video examples of Tk-GAL4^FruM^ → CsChrimson, *R82C10-LexA* → Shibire^TS^ flies fighting at 20°C (permissive temperature) and 30°C (restrictive temperature) during the final LED stimulation period. Pairs in 20°C formed winner-loser hierarchy by this time, while pairs in 30°C continued engaging in active fights. The episodes start 5s before the onset of 6^th^ LED stimulation.

Supplemental Video 4: Blocking the synaptic transmission of V2 MBONs disrupts defeat onset, related to Figure 4.

Video examples of Tk-GAL4^FruM^ → CsChrimson, *R65B09-LexA* → Shibire^TS^ flies fighting at 20°C (permissive temperature) and 30°C (restrictive temperature) during the final LED stimulation period. Pairs in 20°C formed winner-loser hierarchy by this time, while pairs in 30°C continued engaging in active fights. The episodes start 5s before the onset of 6^th^ LED stimulation.

---

**Supplemental Figure S1:**
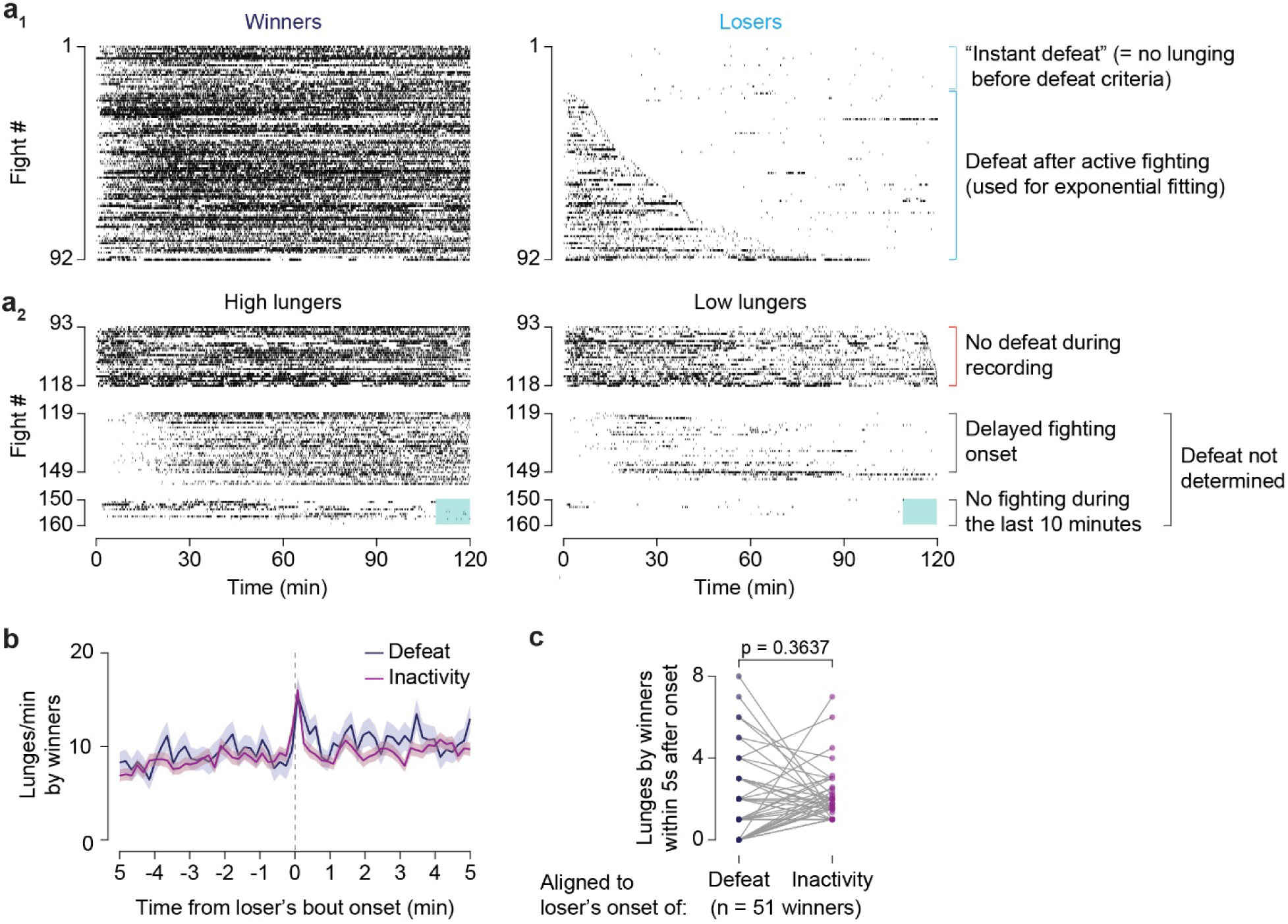
Dynamics of fighting and dominance of wild-type males. **a)** Breakdown of all fights between socially isolated Canton-S males collected in this study, separated into pairs in which defeat was determined (**a_1_**, data set for Fig. 1a) and pairs in which defeat was either absent or undetermined (**a_2_**). Lunge rasters sorted by loser lunge activity. **a_1_)** Yellow bracket: fights in which the loser did not fight back while being attacked by the winner during the first 10min of the recording (instant defeat). Instant defeat time is classified as t = 0min (see Methods). These flies are not included in fitting the rate of defeat onset (Fig. 1f). Green bracket: fights in which the loser fought before later succumbing to defeat. **a_2_)** Top raster: Fights with no loser during the duration of the recording. Middle raster: Fights that exhibited significant delays before fighting occur (see Method for details). Bottom: Pairs that did not fight during the dominance determination window (the last 10 minutes: see Method for details). **b)** Mean winner lunge rate aligned aligned to the defeat onset (n = 69 flies, dark blue), and the rate aligned to pre-defeat inactivity bouts (n = 335 bouts from 58 flies, purple) when the loser did not lunge for at least 1 min. Winner lunge rates were computed in 10-s bins within a ±5-min window around a bout. Pairs analyzed were restricted to defeat/inactivity bouts that occurred between 5 and 100 min into the recording. In both cases, lunge rates transiently increased at bout onset. Shaded regions indicate S.E.M.; solid lines denote mean. **c)** Paired analysis of winner lunge counts during the 5s window following either the moment of defeat or loser inactivity (≥1-min periods without lunges as above). For each loser, the mean number of lunges received across all detected inactivity bouts (see b_1_) was compared with the number observed immediately after defeat (see b_2_). No systematic correlation in winner lunging was observed with defeat onset (paired t-test).

**Supplemental Figure S2.**
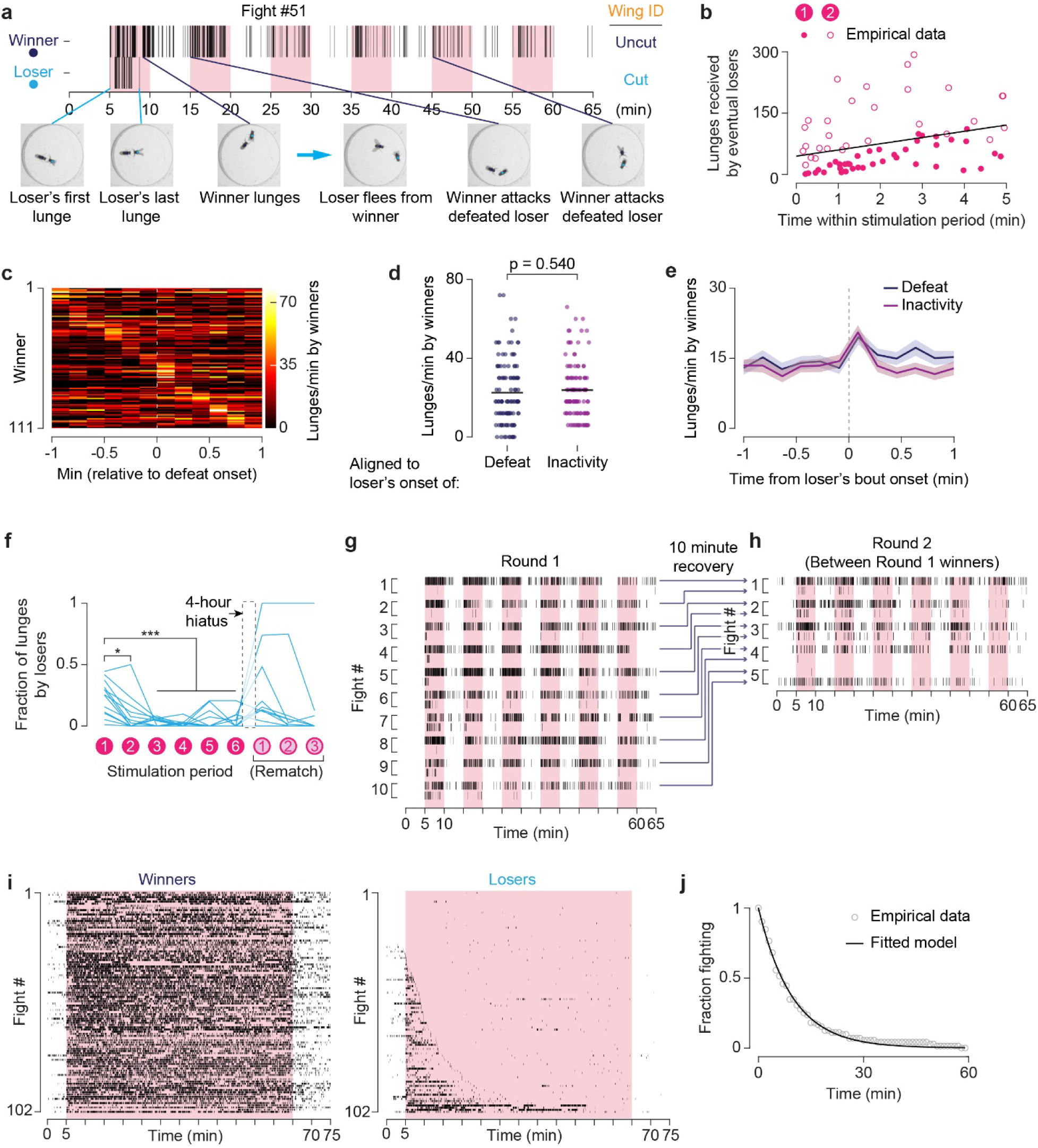
Additional analyses of fights induced by optogenetic activation of Tk-GAL4^FruM^ neurons. **a)** Raster plot of lunges exchanged between an example pair of flies that formed a winner-loser hierarchy. Top: Lunge raster. Bottom: Representative frames from fighting phases that occurred before defeat. **b)** Cumulative number of lunges received by losers plotted against their defeat time within the 1^st^ (closed circles, n = 41) or 2^nd^ (open-circles, n = 29) stimulation period. Compared to losers that were defeated in the 1^st^ stimulus period, losers that were defeated in the 2^nd^ stimulus period received more lunges cumulatively, arguing against a threshold of received lunges before defeat. A solid line is a linear regression of the combined datasets (r = 0.32, p = 0.007). **c)** Heatmap plot of winners’ peak lunge rate (10-s bin) aligned to their opponents defeat onset time (t = 0). Each row is a winner fly, sorted by time of peak lunge rate. Pairs in which the loser was defeated within the first 1-min were excluded, n = 111 winners. **d)** Peak winner lunge rates upon loser defeat (±10-s bin) (n=111, as in **c**) or pre-defeat inactivity bouts when the loser did not lunge for at least 1 min (105 bouts, 42 flies). Welch’s t-test. **e)** Mean winner lunge rates aligned to loser defeat onset (n = 111 flies) or 1-min loser inactivity bouts (as in **d**, n = 105 bouts, from 42 losers). Winner lunge rate transiently increased at both bout onsets to similar levels. Trace shading = S.E.M. **f)** The fraction of lunges by losers relative to total lunges per pair performed during each stimulation period during the fighting paradigm in Fig. 2a, as well as during the rematch with 3 additional stimulation periods after 4-hours of no stimulation. Most flies remained unresponsive to Tk-GAL4^FruM^-neuron activation, but 3 recovered to become the dominant lunger. n = 13 flies, one-way ANOVA, * p < 0.05, *** p < 0.001. **g, h**) Successive fights between winners of fights induced by optogenetic activation of Tk-GAL4^FruM^ neurons. (**g**) Lunge raster of “round 1” fights. Fighting paradigm as in Fig. 2 a. Each fight (bracket #) is sorted winner top, loser bottom. (**h**) Lunge rasters of “Round 2” fights between winners of the Round 1. Each fight (bracket #) is again sorted winner top, loser bottom. **i)** Paired winner-loser lunge rasters for fights induced by optogenetic activation of Tk-GAL4^FruM^ neurons with 1-hr constant stimulation (10ms pulses at 5Hz). Flies sorted by defeat time. **j)** Proportion of losers still fighting during stimulation (circles, 1-min bins), fit by an exponential function (τ = 10.33 min, 95% CI = 10.01–10.66), n = 75 losers (excludes “instant defeat” flies). All panels: pink bars indicate stimulation periods.

**Supplemental Figure S3.**
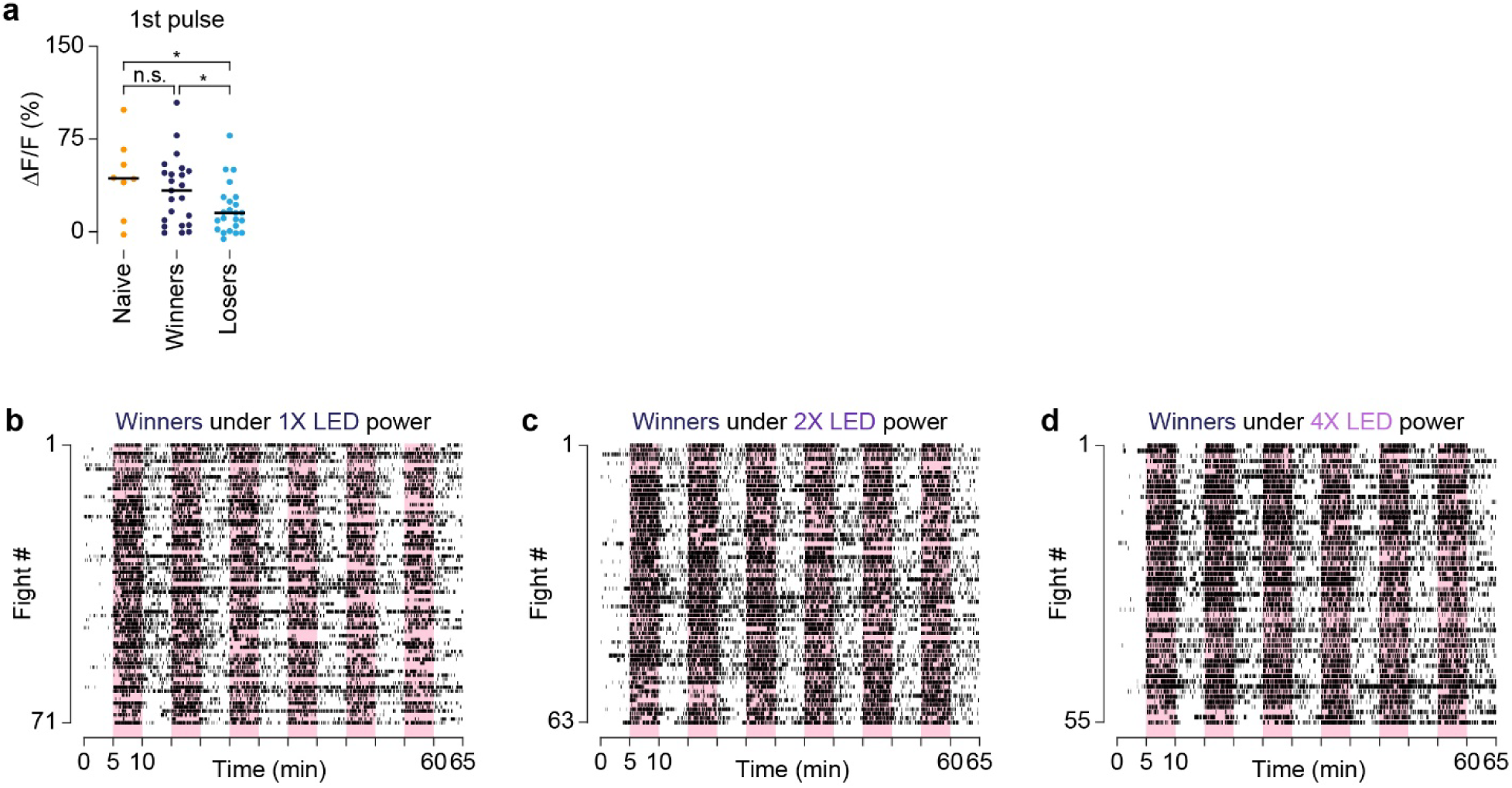
Supporting analysis of Tk-GAL4^FruM^ neuronal activity. (**a**) Peak GCaMP7s responses of Tk-GAL4^FruM^ neurons to optogenetic activation during the first stimulation periods of naive (no opponent in fighting chamber; orange, n = 8), winners (dark blue, n = 23) and losers (cyan, n = 23); n.s. p > 0.05, * p < 0.05 by unpaired t-test. (**b-d**) Lunge rasters of winners using 1X, 2X, and 4X LED stimulation intensities corresponding to losers in Fig. 3g-i.

**Supplemental Figure S4.**
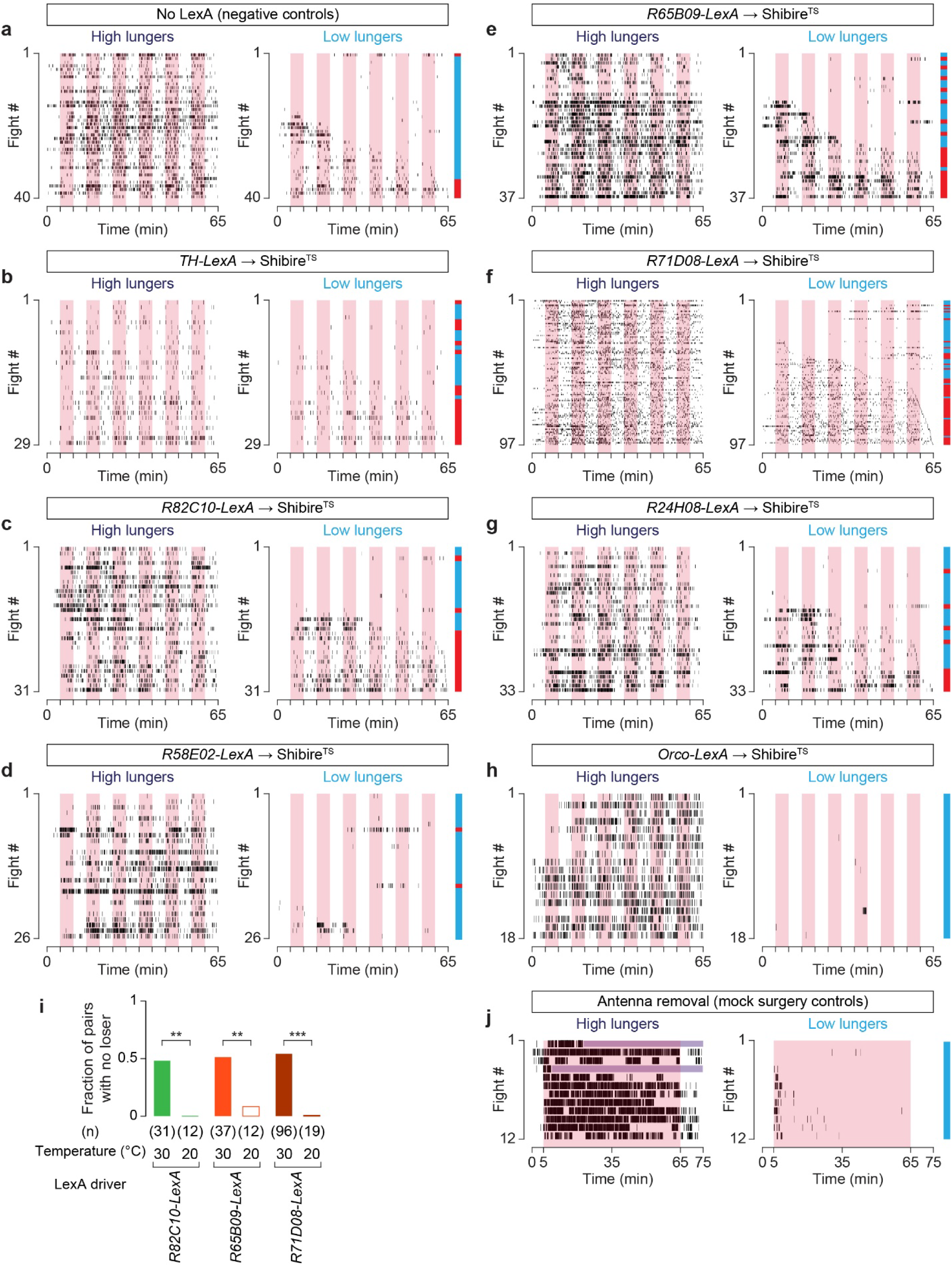
Fighting dynamics under the simultaneous stimulation of Tk-GAL4^FruM^ neurons and blockade of synaptic transmission from select neurons. **a-h**) Paired lunge rasters for all fights among Tk-GAL4^FruM^-neurons → CsChrimson, LexA → Shibire^TS^ genoytpes tested, separated into high and low lungers and ordered by the timing of the low-lunger’s first pause in lunging for 10 minutes or longer (same criterion applied for sorting defeat onset). Color side-bar marks dominance hierarchies determined by the integrated facing angle and lunging dominance classifier (see Methods). Blue-side bar marks pairs that formed winner-loser dominance hierarchies, red bars indicate pairs that didn’t form dominance. **i)** Fraction of fights among Tk-GAL4^FruM^-neurons → CsChrimson, *LexA* → Shibire^TS^ genotypes that did not result in winner-loser hierarchy at 30°C (restrictive temperature) and 20°C (permissive temperature). Chi-square test. ** p < 0.01, *** p < 0.001. **j)** Winner and loser lunge rasters from fights induced by the optogenetic activation of Tk-GAL4^FruM^ neurons between flies that received mock surgery, as a control for Fig. 4h. Purple shading indicates absence of lunging because the loser climbed the wall of fighting chamber. All panels: pink shaded areas indicate periods of CsChrimson stimulation.

**Supplemental Figure S5.**
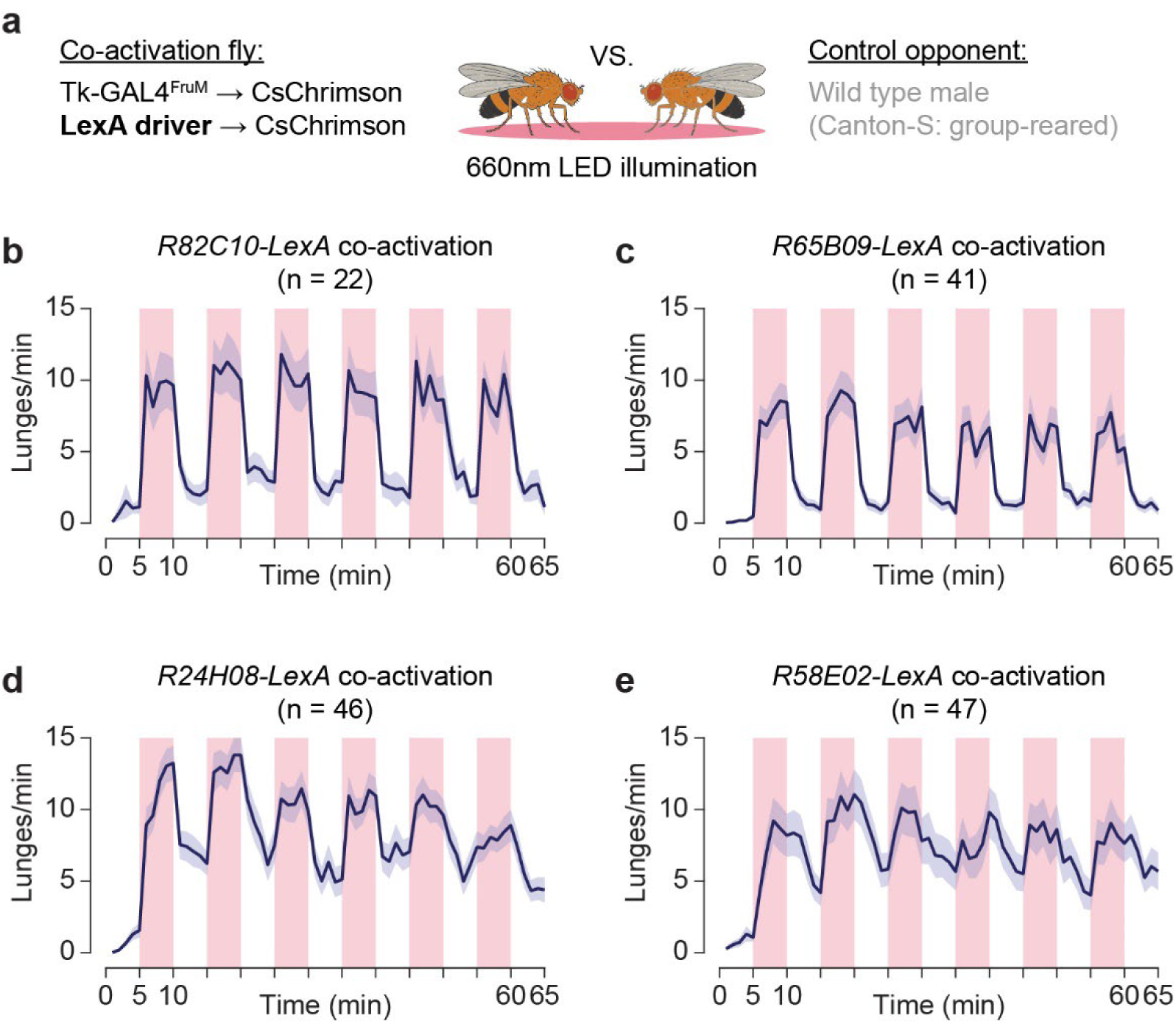
Tk-GAL4^FruM^ neuron activation consistently promotes fighting during co-activation of dopaminergic and MBONs. **a)** Schematic of competition assays designed to test how activating dopaminergic and MBONs affects aggression elicited by the activation of Tk-GAL4^FruM^ neurons. In these assays, flies in which LexA-labeled neurons are co-activated with Tk-GAL4^FruM^ neurons were paired against non-aggressive group-housed Canton-S male flies. **b-e**) Mean lunge rates of co-activation genotypes in response to CsChrimson activation. Pink bars indicate stimulation periods. Trace shading = S.E.M.

**Supplemental Figure S6.**
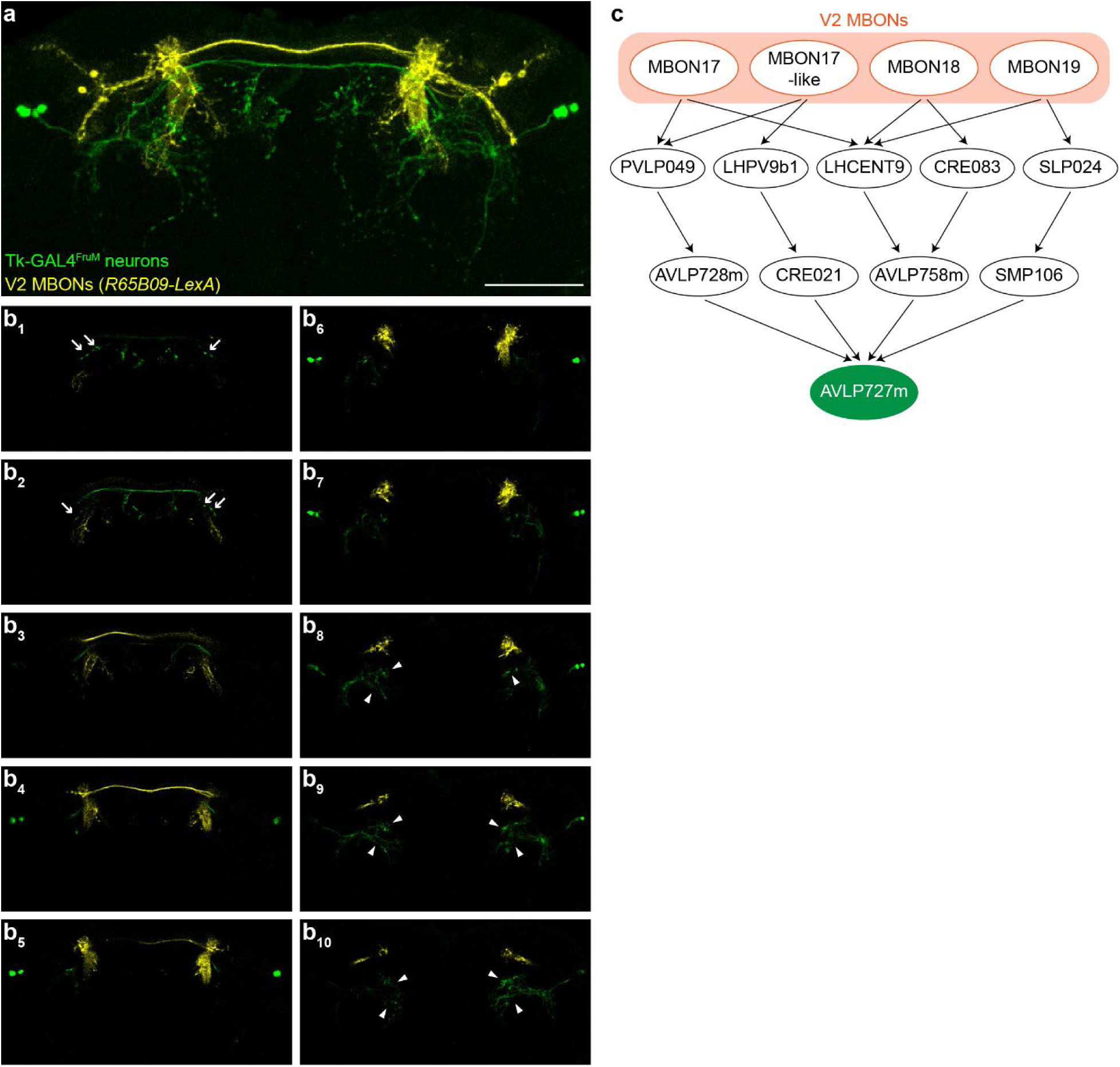
V2 MBONs do not directly synapse with Tk-GAL4^FruM^ neurons. **a)** A representative z-stack of MBONs labeled by *R65B09-LexA* (yellow) and Tk-GAL4^FruM^ neurons (green). MBONs are segmented from other neurons for clarity. Scale bar = 50μm. **b_1-10_**) Serial confocal slices taken every 3μm show that perceived overlaps of V2 MBONs and Tk-GAL4^FruM^ neurons in a) are spatially segregated. Arrows (b_1-2_) indicate putative presynaptic boutons, and white arrowheads (b_8-10_) indicate putative dendritic areas of Tk-GAL4^FruM^ neurons^67^, respectively. **c**) A schematic of the shortest neuronal connectivity paths between V2 MBONs (MBON17-19) and AVLP727m, a cell type annotated to be Tk-GAL4^FruM^ neurons in the male CNS connectome^68^. At least 2 intervening neurons are required to connect V2 MBONs and Tk-GAL4^FruM^ neurons. See Methods for details.

**Supplemental Methods 7.**
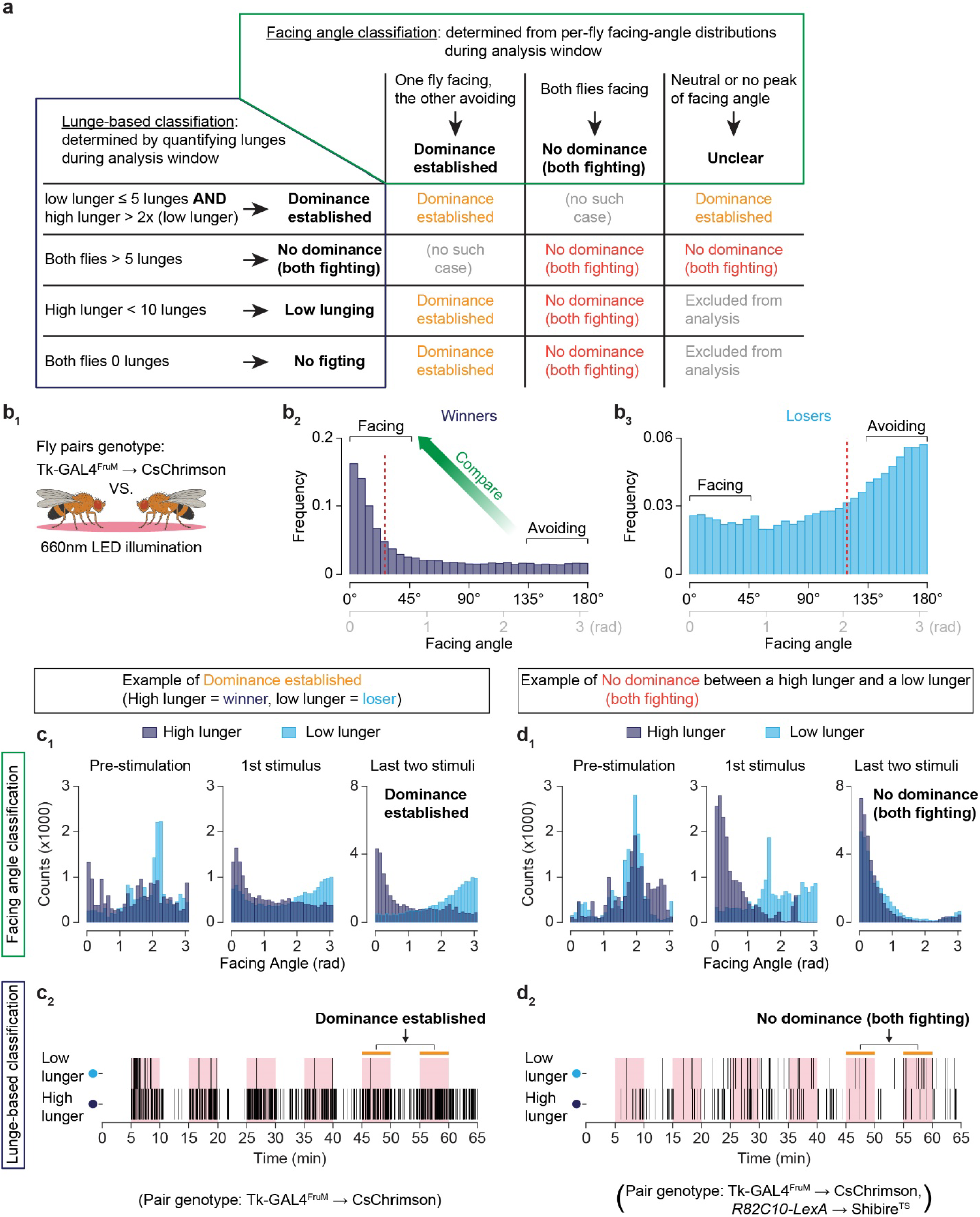
**a)** Overview of the three-stage pipeline used to identify winner–loser dominance: facing-angle classification, lunge-based classification, and a final cross-metric integration step that together confirm when a fly transitions from attacking to fleeing (see Methods). **b)** Facing-angle histograms from Tk-GAL4^FruM^-CsChrimson fights during the dominance-determination window illustrate how orientation is used to classify dominance (data as in Fig. 2c, n = 124 pairs). **(b_1_)** Schematic of assay. **(b_2_)** Winners show a prominent peak in the “facing” range (0–0.75 rad), indicating sustained orientation toward their opponent. **(b_3_)** Losers show a distinct peak in the “fleeing” range (2.4–3.14 rad), indicating orientation away from their opponent. For each fly, the difference between these regions defines its prevailing facing angle, and pairwise comparison of these distributions is used to assign dominance as outlined in (a). Red broken lines indicate median. **c)** Example of a fight classified as “Dominance Established” from a single Tk-GAL4^FruM^-CsChrimson trial. **(c_1_)** Facing-angle histograms and **(c_2_)** lunge rasters are shown for each fly. During the prestimulation period (0–5 min before CsChrimson activation), neither fly showed a facing-angle bias or lunging. During the first stimulation period, Fly 1 (the eventual loser) exhibited a bimodal facing-angle distribution, with peaks in both the “facing” and “fleeing” ranges. This corresponds to its behavioral trajectory: Fly 1 began by initiating attacks, but as stimulation continued, its lunging diminished while Fly 2’s attacks continued. Fly 1 thus showed mixed fighting and fleeing. Fly 2, in contrast, displayed a single dominant peak in its facing-angle distribution—matching the pattern seen in winners—and lunged consistently throughout stimulation. By the time of dominance-classification window, Fly 1’s facing angles consolidated into a single peak in the fleeing range, while Fly 2 maintained a single peak in the facing range. Accordingly, both facing-angle and lunge-based metrics classify the pair as “Dominance Established.” **d)** Example of a fight classified as “No Dominance” from an *R82C10-LexA*→Shibire^TS^ silencing condition, which blocks dominance formation (see main text and Fig. 4). Unlike the example in (c), both flies exhibited peaks in the facing range of the histogram during the dominance-classification window, and both continued lunging. Because neither fly stopped attacking or transitions into a fleeing-biased orientation, the pair remained mutually engaged in fighting, and “no dominance” is assigned. Together, panels **(b–d)** illustrate how combining lunging behavior with facing-angle orientation provides complementary, mutually reinforcing measures of dominance dynamics. For all histograms, counts were collected using 0.1-rad bins spanning 0–π. Pink bars in the raster plots denote CsChrimson stimulation periods.

**Supplementary Table 1:** Complete genotypes used in this study.

| Abbreviated Genotype | Complete Genotype | Figure |
| --- | --- | --- |
| CSMH Wild-type | +/+; +/+; +/+ (Canton-S) | Fig. 1, Supplemental Fig. 1, Fig. 2e |
| *Tk-GAL4<sup>1</sup>→CsChrimson* | *w, Tk-GAL4<sup>1</sup>; Otd-nls:FLPo in attP40/+; 20XUAS>myr:TopHAT2>Syn21-CsChrimson:tdTomato in attP2* | Fig. 2, Supplemental Fig. 2, Fig. 3e-i. Fig. 4h-j. Supplemental Fig. 4j, Fig. 6 |
| *Tk-GAL4<sup>1</sup>→CsChrimson*<br>GCaMP7s<br>Imaging strain | *w, Tk-GAL4<sup>1</sup>; Otd-nls:FLPo in attP40/ 20XUAS-IVS-jGCaMP7s in su(Hw)attP5; 20XUAS>myr:TopHAT2>Syn21-CsChrimson:tdTomato in attP2* | Fig. 3a-d.<br>Supplemental Fig. 3a |
| *Shibire<sup>TS</sup>* - No LexA<br>control<br>*Tk-GAL4<sup>1</sup>→CsChrimson* | *w, Tk-GAL4<sup>1</sup>; Otd-nls:FLPo in attP40 / +; 20xUAS>myr:TopHAT2>CsChrimson:tdTomato in attP2/ 13xLexAop2-IVS-Syn21-shibire<sup>ts</sup> in VK00005* | Fig. 4c, d,<br>Supplemental Fig. 4a |
| *TH-LexA→Shibire<sup>TS</sup>*<br>*Tk-GAL4<sup>1</sup>→CsChrimson* | *w, Tk-GAL4<sup>1</sup>; Otd-nls:FLPo in attP40 / TH-LexA; 20xUAS>myr:TopHAT2>CsChrimson:tdTomato in attP2/ 13xLexAop2-IVS-Syn21-shibire<sup>ts</sup> in VK00005* | Fig. 4c, d,<br>Supplemental Fig. 4b |
| *R82C10-LexA→Shibire<sup>TS</sup>*<br>*Tk-GAL4<sup>1</sup>→CsChrimson* | *w, Tk-GAL4<sup>1</sup>; Otd-nls:FLPo in attP40 / R82C10-LexA in attP40; 20xUAS>myr:TopHAT2>CsChrimson:tdTomato in attP2/ 13xLexAop2-IVS-Syn21-shibire<sup>ts</sup> in VK00005* | Fig. 4c, e,<br>Supplemental Fig. 4c, i |
| *R58E02-LexA→Shibire<sup>TS</sup>*<br>*Tk-GAL4<sup>1</sup>→CsChrimson* | *w, Tk-GAL4<sup>1</sup>; Otd-nls:FLPo in attP40 / R58E02-LexA in attP40; 20xUAS>myr:TopHAT2>CsChrimson:tdTomato in attP2/ 13xLexAop2-IVS-Syn21-shibire<sup>ts</sup> in VK00005* | Fig. 4c, d,<br>Supplemental Fig. 4d |
| *R65B09-LexA→Shibire<sup>TS</sup>*<br>*Tk-GAL4<sup>1</sup>→CsChrimson* | *w, Tk-GAL4<sup>1</sup>; Otd-nls:FLPo in attP40 / R65B09-LexA in attP40; 20xUAS>myr:TopHAT2>CsChrimson:tdTomato in attP2/ 13xLexAop2-IVS-Syn21-shibire<sup>ts</sup> in VK00005* | Fig. 4c, f,<br>Supplemental Fig. 4e, i |
| *R71D08-LexA→Shibire<sup>TS</sup>*<br>*Tk-GAL4<sup>1</sup>→CsChrimson* | *w, Tk-GAL4<sup>1</sup>; Otd-nls:FLPo in attP40 / R71D08-LexA in attP40; 20xUAS>myr:TopHAT2>CsChrimson:tdTomato in attP2/ 13xLexAop2-IVS-Syn21-shibire<sup>ts</sup> in VK00005* | Fig. 4c, f,<br>Supplemental Fig. 4f, i |
| *R24H08-LexA→Shibire<sup>TS</sup>*<br>*Tk-GAL4<sup>1</sup>→CsChrimson* | *w, Tk-GAL4<sup>1</sup>; Otd-nls:FLPo in attP40 / R24H08-LexA in attP40; 20xUAS>myr:TopHAT2>CsChrimson:tdTomato in attP2/ 13xLexAop2-IVS-Syn21-shibire<sup>ts</sup> in VK00005* | Fig. 4b.<br>Supplemental Fig. 4g. |
| *Orco-LexA→Shibire<sup>TS</sup>*<br>*Tk-GAL4<sup>1</sup>→CsChrimson* | *w, Tk-GAL4<sup>1</sup>; Otd-nls:FLPo in attP40 / 20xUAS>myr:TopHAT2>CsChrimson:tdTomato in VK000022; Orco-LexA/13xLexAop2-IVS-Syn21-shibire<sup>ts</sup> in VK00005* | Fig. 4b.<br>Supplemental Fig. 4h. |
| *R82C10-LexA→CsChrimson*<br>*Tk-GAL4<sup>1</sup>→CsChrimson* | *w, Tk-GAL4<sup>1</sup>; Otd-nls:FLPo in attP40 / R82C10-LexA in attP40; 20xUAS>myr:TopHAT2>CsChrimson:tdTomato in attP2/ 13xLexAop2-IVS-Syn21-CsChrimson:tdTomato in attP2* | Fig. 5b, f-g |
| *R65B09-LexA→CsChrimson*<br>*Tk-GAL4<sup>1</sup>→CsChrimson* | *w, Tk-GAL4<sup>1</sup>; Otd-nls:FLPo in attP40 / R65B09-LexA in attP40; 20xUAS>myr:TopHAT2>CsChrimson:tdTomato in* | Fig. 5c, f-g |
|  | *attP2/ 13xLexAop2-IVS-Syn21-CsChrimson:tdTomato* in *attP2* |  |
| *R24H08-LexA→CsChrimson*<br>*Tk-GAL4<sup>1</sup>→CsChrimson* | *w, Tk-GAL4<sup>1</sup>; Otd-nls:FLPo* in *attP40* / *R24H08-LexA* in *attP40*;<br>*20xUAS>myr:TopHAT2>CsChrimson:tdTomato* in *attP2/ 13xLexAop2-IVS-Syn21-CsChrimson:tdTomato* in *attP2* | Fig. 5d, f-g |
| *R58E02-LexA→CsChrimson*<br>*Tk-GAL4<sup>1</sup>→CsChrimson* | *w, Tk-GAL4<sup>1</sup>; Otd-nls:FLPo* in *attP40* / *R58E02-LexA* in *attP40*;<br>*20xUAS>myr:TopHAT2>CsChrimson:tdTomato* in *attP2/ 13xLexAop2-IVS-Syn21-CsChrimson:tdTomato* in *attP2* | Fig. 5e, f-g |
| *R65B09-LexA and Tk-GAL4<sup>FruM</sup> neurons* | *w, Tk-GAL4<sup>1</sup>; R65B09-LexA* in *attP40* / *20xUAS>myr:TopHAT2>GCaMP6f* in *su(Hw)attP5; fru<sup>FLP</sup> / 13xLexAop2-IVS-Syn21-CsChrimson:tdTomato* in *attP2* | Supplemental Fig. 6a |

## References

1. Huntingford, F. & Turner, A. K. Animal Conflict. (Chapman and Hall, London; New York, 1987).

2. Maynard Smith, J. Evolution and the Theory of Games. (Cambridge University Press, Cambridge ; New York, 1982).

3. Hardy, I. C. W. & Briffa, M. Animal Contests. (Cambridge University Press, 2013).

4. Pereira, T. D., Shaevitz, J. W. & Murthy, M. Quantifying behavior to understand the brain. Nat Neurosci 23, 1537–1549 (2020).

5. Venken, K. J. T., Simpson, J. H. & Bellen, H. J. Genetic manipulation of genes and cells in the nervous system of the fruit fly. Neuron 72, 202–230 (2011).

6. Chen, S., Lee, A. Y., Bowens, N. M., Huber, R. & Kravitz, E. A. Fighting fruit flies: A model system for the study of aggression. Proc Natl Acad Sci USA 99, 5664–5668 (2002).

7. Asahina, K. Neuromodulation and strategic action choice in *Drosophila* aggression. *Ann*. Rev. Neurosci. 40, 51–75 (2017).

8. Yurkovic, A., Wang, O., Basu, A. C. & Kravitz, E. A. Learning and memory associated with aggression in *Drosophila melanogaster*. Proc Natl Acad Sci USA 103, 17519–17524 (2006).

9. Trannoy, S., Penn, J., Lucey, K., Popovic, D. & Kravitz, E. A. Short and long-lasting behavioral consequences of agonistic encounters between male *Drosophila melanogaster*. Proc Natl Acad Sci U S A 113, 4818–4823 (2016).

10. Hu, S. W., Yang, Y. T., Sun, Y., Zhan, Y. P. & Zhu, Y. Serotonin signals overcome loser mentality in *Drosophila*. iScience 23, 101651 (2020).

11. Simon, J. C. & Heberlein, U. Social hierarchy is established and maintained with distinct acts of aggression in male Drosophila melanogaster. J. Exp. Biol. 223, (2020).

12. Asahina, K. Neuromodulation of conflicts and hierarchy in insects. in Comprehensive Molecular Insect Science (Second Edition) (eds Yamanaka, N. & Atkinson, P. W.) 263–280 (Elsevier, Oxford, 2026).

13. Auer, T. O. & Benton, R. Sexual circuitry in *Drosophila*. Curr Opin Neurobiol 38, 18–26 (2016).

14. Asahina, K. et al. Tachykinin-expressing neurons control male-specific aggressive arousal in *Drosophila*. Cell 156, 221–235 (2014).

15. Chiu, H. et al. A circuit logic for sexually shared and dimorphic aggressive behaviors in *Drosophila*. Cell 184, 507–520.e16 (2021).

16. Hoopfer, E. D., Jung, Y., Inagaki, H. K., Rubin, G. M. & Anderson, D. J. P1 interneurons promote a persistent internal state that enhances inter-male aggression in *Drosophila*. eLife 4, e11346 (2015).

17. Wohl, M., Ishii, K. & Asahina, K. Layered roles of *fruitless* isoforms in specification and function of male aggression-promoting neurons in *Drosophila*. eLife 9, e52702 (2020).

18. Kim, Y.-K. et al. Repetitive aggressive encounters generate a long-lasting internal state in *Drosophila melanogaster* males. Proc Natl Acad Sci USA 115, 1099–1104 (2018).

19. Klapoetke, N. C. et al. Independent optical excitation of distinct neural populations. Nat Methods 11, 338–346 (2014).

20. Dana, H. et al. High-performance calcium sensors for imaging activity in neuronal populations and microcompartments. Nat Methods 16, 649–657 (2019).

21. Wohl, M. P., Liu, J. & Asahina, K. *Drosophila* tachykininergic neurons modulate the activity of two groups of receptor-expressing neurons to regulate aggressive tone. J Neurosci 43, 3394–3420 (2023).

22. Kitamoto, T. Conditional modification of behavior in *Drosophila* by targeted expression of a temperature-sensitive shibire allele in defined neurons. J Neurobiol 47, 81–92 (2001).

23. Galili, D. S. et al. Converging circuits mediate temperature and shock aversive olfactory conditioning in *Drosophila*. Curr Biol 24, 1712–1722 (2014).

24. Hattori, D. et al. Representations of novelty and familiarity in a mushroom body compartment. Cell 169, 956–969.e17 (2017).

25. Liu, C. et al. A subset of dopamine neurons signals reward for odour memory in *Drosophila*. Nature 488, 512–516 (2012).

26. Aso, Y. et al. Mushroom body output neurons encode valence and guide memory-based action selection in *Drosophila*. eLife 3, e04580 (2014).

27. Séjourné, J. et al. Mushroom body efferent neurons responsible for aversive olfactory memory retrieval in *Drosophila*. Nat Neurosci 14, 903–910 (2011).

28. Felsenberg, J., Barnstedt, O., Cognigni, P., Lin, S. & Waddell, S. Re-evaluation of learned information in *Drosophila*. Nature 544, 240–244 (2017).

29. Aso, Y. et al. The neuronal architecture of the mushroom body provides a logic for associative learning. eLife 3, e04577 (2014).

30. Larsson, M. C., et al. *Or83b* encodes a broadly expressed odorant receptor essential for *Drosophila* olfaction. Neuron 43, 703–714 (2004).

31. Task, D. et al. Chemoreceptor co-expression in *Drosophila melanogaster* olfactory neurons. eLife 11, e72599 (2022).

32. Hoffmann, A. A. & Cacoyianni, Z. Territoriality in *Drosophila melanogaster* as a conditional strategy. Anim. Behav. 40, 526–537 (1990).

33. Lim, R. S., Eyjólfsdóttir, E., Shin, E., Perona, P. & Anderson, D. J. How food controls aggression in *Drosophila*. PLoS ONE 9, e105626 (2014).

34. Ueda, A. & Kidokoro, Y. Aggressive behaviours of female *Drosophila melanogaster* are influenced by their social experience and food resources. Physiol. Entomol. 27, 21–28 (2002).

35. Steck, K. et al. Internal amino acid state modulates yeast taste neurons to support protein homeostasis in *Drosophila*. eLife 7, e31625 (2018).

36. Ribeiro, C. & Dickson, B. J. Sex peptide receptor and neuronal TOR/S6K signaling modulate nutrient balancing in *Drosophila*. Curr Biol 20, 1000–1005 (2010).

37. Dow, M. A. & Schilcher, F. V. Aggression and mating success in Drosophila melanogaster. Nature 254, 511–512 (1975).

38. Sturtevant, A. H. Experiments on sex recognition and the problem of sexual selection in *Drosoophilia*. *J*. Anim. Behav. 5, 351–366 (1915).

39. Suárez-Grimalt, R., Grunwald Kadow, I. C. & Scheunemann, L. An integrative sensor of body states: how the mushroom body modulates behavior depending on physiological context. Learn. Mem. 31, a053918 (2024).

40. Keleman, K. et al. Dopamine neurons modulate pheromone responses in *Drosophila* courtship learning. Nature 489, 145–149 (2012).

41. Zhao, X., Lenek, D., Dag, U., Dickson, B. J. & Keleman, K. Persistent activity in a recurrent circuit underlies courtship memory in *Drosophila*. eLife 7, e31425 (2018).

42. Cohn, R., Morantte, I. & Ruta, V. Coordinated and compartmentalized neuromodulation shapes sensory processing in *Drosophila*. Cell 163, 1742–1755 (2015).

43. Siju, K. P. et al. Valence and state-dependent population coding in dopaminergic neurons in the fly mushroom body. Curr Biol 30, 2104–2115.e4 (2020).

44. Zolin, A. et al. Context-dependent representations of movement in *Drosophila* dopaminergic reinforcement pathways. Nat Neurosci 24, 1555–1566 (2021).

45. Aimon, S. et al. Fast near-whole–brain imaging in adult *Drosophila* during responses to stimuli and behavior. PLoS Biology 17, e2006732 (2019).

46. Villar, M. E. et al. Differential coding of absolute and relative aversive value in the *Drosophila* brain. Curr Biol 32, 4576–4592.e5 (2022).

47. Hige, T., Aso, Y., Modi, M. N., Rubin, G. M. & Turner, G. C. Heterosynaptic plasticity underlies aversive olfactory learning in *Drosophila*. Neuron 88, 985–998 (2015).

48. Adel, M. & Griffith, L. C. The role of dopamine in associative learning in *Drosophila*: an updated unified model. Neurosci. Bull. 37, 831–852 (2021).

49. Takemura, S. et al. A connectome of a learning and memory center in the adult *Drosophila*brain. eLife 6, e26975 (2017).

50. Alekseyenko, O. V., Chan, Y.-B., Li, R. & Kravitz, E. A. Single dopaminergic neurons that modulate aggression in *Drosophila*. Proc Natl Acad Sci USA 110, 6151–6156 (2013).

51. Hsu, Y., Earley, R. L. & Wolf, L. L. Modulation of aggressive behaviour by fighting experience: mechanisms and contest outcomes. Biol. Rev. 81, 33–74 (2006).

52. Hammels, C. et al. Defeat stress in rodents: From behavior to molecules. Neurosci Biobehav Rev 59, 111–140 (2015).

53. Diaz, V. & Lin, D. Neural circuits for coping with social defeat. Curr Opin Neurobiol 60, 99–107 (2020).

54. Stevenson, P. A. & Schildberger, K. Mechanisms of experience dependent control of aggression in crickets. Curr Opin Neurobiol 23, 318–323 (2013).

55. Chou, M.-Y. et al. Social conflict resolution regulated by two dorsal habenular subregions in zebrafish. Science 352, 87–90 (2016).

56. Nakajo, H. et al. Hunger potentiates the habenular winner pathway for social conflict by orexin-promoted biased alternative splicing of the AMPA receptor gene. Cell Rep 31, 107790 (2020).

57. Li, H., Zhao, Z., Jiang, S. & Wu, H. Brain circuits that regulate social behavior. Mol Psychiatry 30, 3240–3256 (2025).

58. Ferreira-Fernandes, E. & Peça, J. The neural circuit architecture of social hierarchy in rodents and primates. Front Cell Neurosci 16, (2022).

59. Zhou, T. et al. History of winning remodels thalamo-PFC circuit to reinforce social dominance. Science 357, 162–168 (2017).

60. Zhang, C. et al. Dynamics of a disinhibitory prefrontal microcircuit in controlling social competition. Neuron 110, 516–531.e6 (2022).

61. Xin, Q. et al. Deconstructing the neural circuit underlying social hierarchy in mice. Neuron 113, 444–459.e7 (2025).

62. Nelson, A. C. et al. Molecular and neural control of social hierarchy by a forebrain-thalamocortical circuit. Cell 188, 5535–5554.e23 (2025).

63. Hillman, K. L. Cost-benefit analysis: the first real rule of fight club? Front Neurosci 7, (2013).

64. Yu, J. Y., Kanai, M. I., Demir, E., Jefferis, G. S. X. E. & Dickson, B. J. Cellular organization of the neural circuit that drives *Drosophila* courtship behavior. Curr Biol 20, 1602–1614 (2010).

65. Pfeiffer, B. D., Truman, J. W. & Rubin, G. M. Using translational enhancers to increase transgene expression in *Drosophila*. Proc Natl Acad Sci USA 109, 6626–6631 (2012).

66. Lai, S.-L. & Lee, T. Genetic mosaic with dual binary transcriptional systems in *Drosophila*. Nat Neurosci 9, 703–709 (2006).

67. Hoyer, S. C. et al. Octopamine in male aggression of *Drosophila*. Curr Biol 18, 159–167 (2008).

68. Leng, X., Wohl, M., Ishii, K., Nayak, P. & Asahina, K. Quantifying influence of human choice on the automated detection of *Drosophila* behavior by a supervised machine learning algorithm. PLoS One 15, e0241696 (2020).

69. Ishii, K., Wohl, M., DeSouza, A. & Asahina, K. Sex-determining genes distinctly regulate courtship capability and target preference via sexually dimorphic neurons. eLife 9, e52701 (2020).

70. Eyjolfsdottir, E. et al. Detecting social actions of fruit flies. in Computer Vision – ECCV 2014 (eds Fleet, D., Pajdla, T., Schiele, B. & Tuytelaars, T.) 772–787 (Springer International Publishing, Cham, 2014). doi:10.1007/978-3-319-10605-2_50.

71. Kabra, M., Robie, A. A., Rivera-Alba, M., Branson, S. & Branson, K. JAABA: interactive machine learning for automatic annotation of animal behavior. Nat Methods 10, 64–67 (2013).

72. Wang, J. W., Wong, A. M., Flores, J., Vosshall, L. B. & Axel, R. Two-photon calcium imaging reveals an odor-evoked map of activity in the fly brain. Cell 112, 271–282 (2003).

73. Schindelin, J., et al. Fiji: an open-source platform for biological-image analysis. Nat Methods 9, 676–682 (2012).

74. Berg, S. et al. Sexual dimorphism in the complete connectome of the *Drosophila* male central nervous system. bioRxiv 2025.10.09.680999 (2025) doi:10.1101/2025.10.09.680999.

